# Fitness-valley crossing in subdivided asexual populations

**DOI:** 10.1101/079624

**Authors:** Michael R. McLaren

## Abstract

Adaptations may require multiple mutations that are beneficial only in combination. To adapt, a lineage must acquire mutations that are individually neutral or deleterious before gaining the beneficial combination, thereby crossing a plateau or valley, respectively, in the mapping from genotype to fitness. Spatial population structure can facilitate plateau and valley crossing by allowing neutral and deleterious lineages to survive longer and produce more beneficial mutants. Here, we analyze adaptation across a two-mutation plateau or valley in an asexual population that is subdivided into discrete subpopulations, or demes, connected by migration. We describe how subdivision alters the dynamics of adaptation from those in an equally sized unstructured population and give a complete quantitative description of these dynamics for the island migration model. Subdivision can significantly decrease the waiting time for the adaptation if demes and migration rates are small enough that single-mutant lineages fix in one or more demes before producing the beneficial double mutant. But, the potential decrease is small in very large populations and may also be limited by the slow spread of the beneficial mutant in extremely subdivided populations. Subdivision has a smaller effect on the probability that the population adapts very quickly than on the mean time to adapt, which has important consequences in some applications, such as the development of cancer. Our results provide a general and intuitive framework for predicting the effects of spatial structure in other models and in natural populations.

## 1 Introduction

Adaptations that give rise to new biological functions are often *complex*, emerging from the interactions of multiple mutations. Examples include the evolution of aerobic citrate utilization in experimentally evolving populations of *Escherichia coli* (Blount *et al*. 2012) and the evolution in vertebrates of a ligand-activated transcription factor to bind a new target ligand (Ortlund *et al*. 2007) or new target DNA sequence (McKeown *et al*. 2014). The acquisition of drug resistance in pathogenic bacteria (Wang et al. 2002; Weinreich *et al*. 2006) and immune system escape in influenza (Gong *et al*. 2013) and HIV (Silva *et al*. 2010) can also be complex adaptations.

The mutations involved in complex adaptations have effects on function and fitness that depend on which other mutations are present. As a result, some or all of the sequences of mutations that lead from the ancestral to the adapted genotype may have steps that are neutral or deleterious. Such sequences are said to cross a plateau or valley, respectively, in the map from genotype to fitness. Determining how a species can adapt via such a sequence of mutations (in both cases generically referred to as *valley crossing*) is crucial to understanding how and when complex adaptations evolve.

Valley crossing also has implications for the evolution of human pathogens and in the development of cancer. Persistence of drug resistance in bacteria and viruses has been explained by the observation that, in the absence of drugs, drug-resistant and drug-sensitive strains are separated by a fitness valley, making reversion to drug sensitivity difficult (Levin *et al*. 2000; Trindade *et al*. 2009). The evolution of Human Immunodeficiency Virus-1 (HIV-1) to use an alternate host co-receptor, seen in roughly one-half of HIV-1 patients, may also require crossing a valley (Regoes and Bonhoeffer 2005; Silva *et al*. 2010; Silva and Wyatt 2014). Finally, cancer usually results from a sequence of mutations (often called *hits*) accumulating within an individual cell that give the cell’s lineage an increased growth rate. For example, loss of function (LOF) mutations may be needed in both copies of a particular tumor-suppressor gene (TSG) before a lineage begins growing into a tumor (Knudson 2001; Frank 2007). If both LOF mutations are needed for cells to have a competitive advantage over cells with no mutations, then inactivating the TSG requires crossing a valley or plateau.

In asexual populations, valley crossing requires a clonal lineage to sequentially acquire all of the necessary mutations. Large asexual populations typically cross plateaus and valleys by a process in which a lineage acquires the mutations while still rare in the population. This process also occurs in sexual populations if the mutations are closely linked (Weissman et al. 2010), as is likely if the mutations occur in the same gene. Komarova *et al*. (2003) used the term *tunneling* to distinguish this process from the typical mode of adaptation in small populations by which the mutations fix sequentially. They were also the first to analyze tunneling where two mutations are required and the single-mutant genotype is either neutral or deleterious. Gillespie (1984) first analyzed tunneling for strongly deleterious intermediate mutants and Kimura (1985) analyzed tunneling as a form of neutral evolution, where the single mutant is deleterious and the double mutant is neutral. Later studies analyzed asexual tunneling under new regimes or provided additional details (e.g., Iwasa *et al*. 2004, Weinreich and Chao 2005, Durrett and Schmidt 2008, and Proulx 2011; see Weissman *et al*. (2009) for the most complete account of valley crossing and tunneling in asexual populations). Genomic evidence suggests that tunneling occurred repeatedly in the evolution of mitochondrial tRNA genes (Meer *et al*. 2010). Theory, observations, and experiments suggest that tunneling occurs during co-receptor switch in HIV (Regoes and Bonhoeffer 2005; Silva and Wyatt 2014). The large size of stem cell populations also suggests that tunneling will often be involved in the acquisition of multiple hits leading to some cancers. Even in cases where intermediate genotypes offer a small fitness advantage, tunneling may be a more likely mode of adaptation that sequential fixation of individual mutations (Weissman *et al*. 2009).

Tunneling studies typically assume that the population is completely unstructured, with all individuals competing equally with one another; however, natural populations are spatially structured to various degrees, such that individuals disperse and compete within a region smaller than the population’s total range. Spatially structured populations can be subdivided into distinct subpopulations, or *demes*; distributed continuously in space; or fall somewhere in between these extremes. Early on, Sewall Wright suggested that spatial structure should facilitate valley crossing (Wright 1932). Wright was mainly concerned with sexual species having high rates of recombination between loci and, in these species, spatial structure reduces the rate at which sex breaks up beneficial combinations of mutations. This idea forms the backbone of Wright’s famous shifting balance theory of evolution (Wright 1970; Charlesworth and Charlesworth 2010). However, the extensive literature on the shifting balance theory says little about how spatial structure might affect valley crossing in asexual species.

We aim to understand how spatial structure affects valley crossing in asexual populations for the simplest scenario, where the adaptation requires two mutations. Spatial structure can have qualitatively different effects on evolutionary dynamics depending on assumptions about how competition and demography interact with space. We set out to describe the consequences of perhaps the most basic aspect of spatial structure, that individuals tend to compete and reproduce within a local region that is small compared to the population’s total geographic range. This basic feature of space can affect asexual tunneling in the following way. If migration is sufficiently limited, mutant lineages that segregate in the population will tend to be unevenly spatially distributed, and mutant individuals will tend to have a greater than average proportion of mutants nearby. As a result, mutants tend to compete more with other mutants than they would if the population were unstructured. This phenomenon, which we call *positive competitive assortment*, reduces the ability for selection and drift to purge deleterious and neutral mutant lineages from the population. Single-mutant lineages are therefore more likely to survive long enough to successfully tunnel across the valley by producing a double mutant than in unstructured populations.

Komarova (2006) showed that spatial structure can accelerate tunneling across valleys and plateaus by this mechanism. Structure was modeled in the form of a population where each each individual occupies a unique site on a one-dimensional lattice, competes with its nearest neighbors, disperses to adjacent sites, and does not migrate. In this model, single-mutant lineages form contiguous colonies that change slowly in size since genetic change occurs only at the colony’s two edges. As a result, single-mutant lineages survive longer and produce more double mutants, which can lead to faster tunneling on average. In contrast, Takahasi (2007) found no effect of population subdivision on plateau crossing in the absence of recombination; however, their mutation model differs from that of Komarova (2006) by allowing each mutation to occur only once in the history of the population.

The results of Komarova (2006) show that structure can in principle accelerate valley crossing; however, questions remain about when this effect will be significant, which we address in the present study. First, how do the effects of spatial structure on valley crossing vary with the type of structure (e.g., subdivided versus continuous habitats) and the degree of structure (e.g., migration rates)? Second, do limits exist for the extent to which spatial structure can accelerate valley crossing? In other words, are there situations where increasing the degree of structure does not decrease the waiting time for the population to adapt?

For certain applications, the rare times when the valley is crossed very quickly are of primary importance; examples include the development of cancer within a human lifetime (Weissman *et al*. 2010) or if crossing the valley before a mutually exclusive beneficial mutation occurs could lead the population down a divergent long-term evolutionary trajectory. In unstructured populations, the dynamics associated with quickly crossing the valley differ in important ways from those associated with crossing the valley in a roughly average amount of time (Weissman *et al*. 2010). Therefore, we also ask how spatial structure affects the probability that the population crosses the valley within a period of time that is much shorter than the average.

To address these questions, we analyzed how population structure affects the dynamics of adaptation across a two-mutation valley or plateau in an asexual population that is subdivided into discrete subpopulations, or *demes*, with occasional migration between demes. Subdivision may have particularly large effects on tunneling, since a rare lineage that becomes fixed within a deme only competes with itself and not with other genotypes. Subdivision is also relevant for the evolution of pathogens and cancer development, as human tissues create subdivided habitats. The population of stem cells from which colon cancers arise, for example, are subdivided into ~ 10^7^ crypts (Tomasetti and Vogelstein 2015), each with an independently evolving subpopulation of ~ 5 stem cells that are susceptible to developing into cancer (Baker *et al*. 2014; Vermeulen and Snippert 2014), and populations of cells carrying HIV are subdivided into pulps within the spleens of infected patients (Frost *et al*. 2001).

We first describe the general mechanism by which subdivision affects valley crossing and then give a complete description of valley-crossing dynamics for the finite island model of subdivision, in which migration between any pair of demes occurs equally frequently. We consider the effects of varying the degree of spatial structure in the island model by varying the size of demes or the migration rates between them. Though the island model is unrealistic for many species, many of our results are general and provide fundamental insights into how subdivision, and spatial structure more generally, affect valley crossing. Section 7 considers how the results, methods, and intuition we present can be extended to models where migration occurs primarily between nearby demes.

## 2 Model

We consider the process by which a subdivided asexual population obtains an adaptation that requires two mutations before any fitness benefit is gained. The mutations leading to such adaptations can be of a variety of types. For example, two point mutations might be required in the same or different genes. Alternatively, for the case of inactivation of a tumor-suppressor gene in a clonally reproducing diploid cell, a nonsense or frameshift mutation may inactivate one copy of the gene and be followed by gene conversion to inactivate the second copy. We analyze how subdivision changes how the population adapts by a particular sequence of two mutations while ignoring the possibility of other mutations. This assumption simplifies our analysis by allowing us to consider just three types of individuals-wild-type individuals that carry zero mutations and individuals carrying either one or two mutations. Wild types, single mutants, and double mutants have relative fitnesses of 1, 1 − *δ*, and 1 + *s*, respectively, with *δ* ≤ 0 and *s* > 0 (Figure 1). Wild types mutate to single mutants at rate *µ*_0_ and single mutants mutate to double mutants at rate *µ*_1_; for simplicity, we ignore back mutations. In general, a population can adapt by multiple mutational pathways; valley-crossing dynamics can then be found from our results using the method described in Weissman *et al*. (2009).

**Figure 1:**
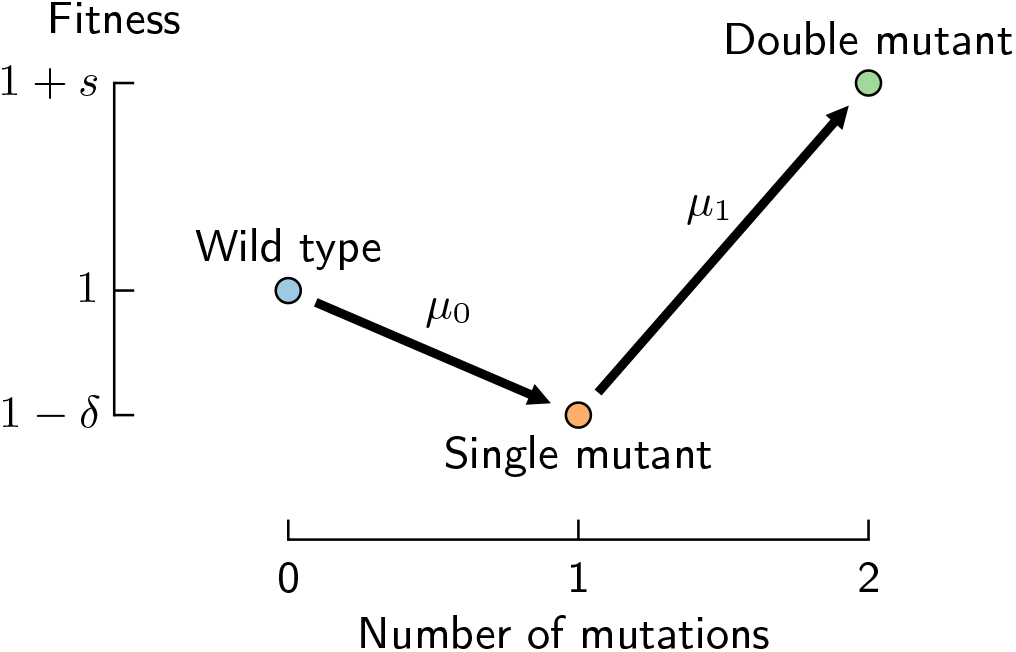
Genotype-to-fitness mapping for a two-mutation fitness valley (*δ* > 0) or plateau (*δ* = 0).

The population consists of *N*_T_ individuals subdivided into *L* demes, with each individual competing and reproducing locally, that is, within its own deme. The number of individuals within a deme is constant and independent of the average fitness of the deme. Migration exchanges individuals between demes but does not alter deme size or the total frequency of genotypes within the population; thus, in our model selection is *soft* and migration is *conservative* (Nagylaki 1980; Whitlock 2004; Charlesworth and Charlesworth 2010). With these assumptions, subdivision affects tunneling by the same general mechanism as in the lattice model of Komarova (2006). We describe how subdivision qualitatively affects valley crossing under a general model where demes may differ in size and allowing for any pattern of migration, so long as each deme can be reached from any other in a finite time. Our quantitative analysis assumes the finite island model, for which each deme has *N* individuals that each migrate at rate *m* per generation to a different, randomly chosen deme.

The rate of genetic drift is controlled by a parameter, *α*, which we call the *drift coefficient*, defined as one-half of the variance in the number of descendants left by an individual after one generation. This definition is equivalent to *α* ≡ *N*/2*N*_e_, where *N*_e_ is the variance effective population size of a deme of size *N*. Larger values of *α* correspond to smaller effective population sizes and faster stochastic changes in genotype frequencies with a deme for a given *N*. Our inclusion of the parameter *α* serves two purposes. First, in many cases the effects of subdivision on the evolution of a mutant lineage can be understood as decreasing the effective drift and selection coefficients. Second, our model can represent a variety of reproduction models by appropriately choosing *α*; for example, the Moran reproduction model corresponds to *α* = 1 and the Wright-Fisher model to *α* = 1/2. Table 1 summarizes our model parameters.

**Table 1:** Model parameters.

|  |  |
| --- | --- |
| $L$ | Number of demes |
| $N_T$ | Census population size of the total population |
| $N$ | Census population size of a deme in the island model |
| $m$ | Migration rate in the island model |
| $\alpha$ | Drift parameter |
| $\mu_0$ | Mutation rate of wild type to single mutant |
| $\mu_1$ | Mutation rate of single to double mutant |
| $\delta$ | Selective disadvantage of single mutant |
| $s$ | Selective advantage of double mutant |

We make several additional assumptions in order to facilitate our analysis and discussion. First, we assume that *s* ≫ *α/N*_T_ so that the double mutant has a significant selective advantage over wild type. Second, we assume that *µ*_1_ ≪ max{*δ*, *s*}, which, for reasons described below, is required for spatial structure to accelerate valley crossing. Third, we assume that the dynamics of genotype frequencies within demes can be described using the standard diffusion approximation from population genetics (Ewens 2004). In practice, this assumption requires that the census and effective sizes of demes are large (*N* ≫ 1 and *N/α* ≫ 1), selection has a small effect on an individual’s reproductive success (*δ* ≪ *α* and *s* ≪ *α*), and that third and higher moments of the distribution of the number of descendants per individual per generation are negligible (i.e., the distribution is characterized only by its mean and variance). Finally, unless otherwise stated we assume that the population is large enough that valley crossing occurs by tunneling—that is, single mutants remain rare in the total population until the double mutant begins to sweep.

We verified our predictions using computer simulations of a Wright-Fisher population. Additional details for our simulations are given in Supplementary Appendix A. Source code for simulations and figures is available at https://github.com/mmclaren42/valley-crossing-subdivided.

## 3 Summary of results

Our goal is to find the average and probability distribution of the time 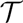 when, starting from an entirely wild-type population, a double mutant first arises whose lineage is destined to sweep through the population, as well as the average time for the double mutant to become fixed within the population.

For the population to adapt across the valley, a single-mutant lineage must survive long enough to produce a double-mutant lineage that is destined to survive and sweep to fixation. We refer to the single and double mutants that give rise to such lineages, and the lineages themselves, as *successful*. Figure 2 illustrates the process of adaptation in a population that is not too large or too heavily subdivided. We let 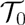 be the waiting time for the first successful single mutant; the *drift time*, 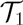, be the time that the first successful single-mutant lineage drifts before producing a successful double mutant; and the *sweep time*, 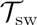, be the time for the successful double-mutant lineage to sweep to fixation. This and other notation is reviewed in Table 2.

**Figure 2:**
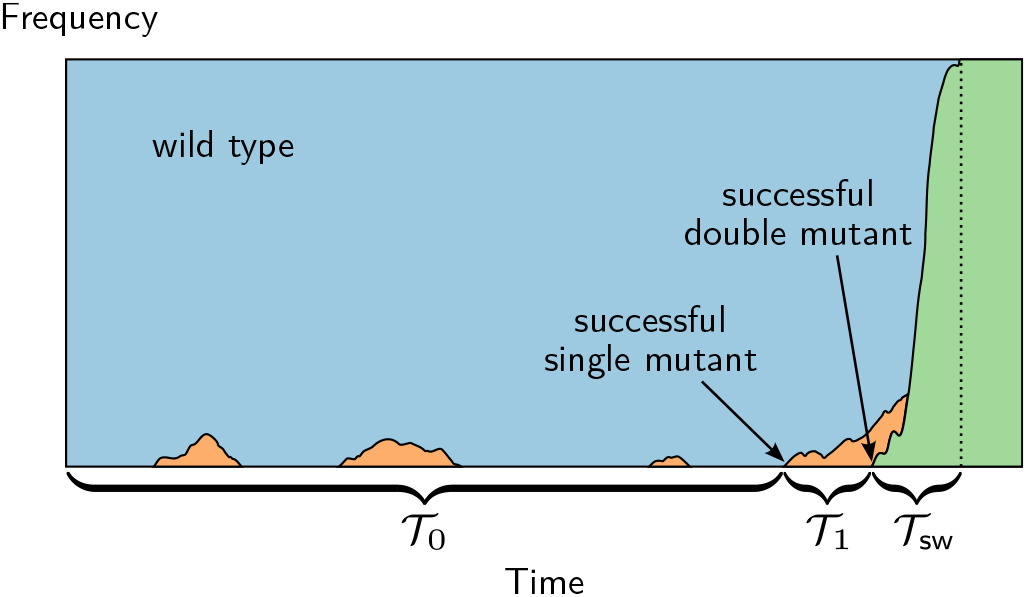
When successful single mutants are infrequent, the time for the population to tunnel across the valley is typically dominated by the time 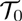 until the first successful single mutant is produced.

**Table 2:** Model parameters.

|  |  |
| --- | --- |
| $p_1$ | Probability that a single mutant is successful |
| $p_2$ | Probability that a double mutant is successful |
| $\mathcal{T}_0$ | Waiting time for the first successful single mutant |
| $\mathcal{T}_1$ | Drift time of the first successful single mutant |
| $\mathcal{T}$ | Waiting time for the first successful double mutant |
| $\mathcal{T}_{\text{sw}}$ | Time for the first successful double mutant to sweep to fixation |
| $p_{1,\text{wm}}$ | Value of $p_1$ in an unstructured population of $N_T$ individuals |
| $\langle \mathcal{T}_{1,\text{wm}} \rangle$ | Value of $\langle \mathcal{T}_1 \rangle$ in an unstructured population of $N_T$ individuals |
| $F$ | Measure of genetic assortment |
| $P_f(i, k)$ | Probability that a mutant lineage with fitness $1 + f$ present in $i$ copies reaches $k$ copies before extinction |
| $\psi_{i \rightarrow j}$ | Probability that a lineage with $j$ mutations fixes in a deme of individuals with $i$ mutations |
| $\theta$ | Probability that a double mutant is successful after fixing in one deme |
| $N_\times$ | Threshold deme size |
| $m^*$ | Migration boundary for the isolated-demes regime |
| $\eta$ | Compound parameter defined in Equation (23) |
| $\langle \mathcal{T} \rangle_{\text{nsd}}$ | $\langle \mathcal{T} \rangle$ under neutral semi-deterministic tunneling |

In Figure 2, the first successful single-mutant lineage produces the first successful double mutant, which in turn becomes the most recent common ancestor of the population once the double-mutant genotype is fixed. Therefore, the time when the first successful double mutant occurs is 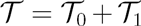 and the time for the population to become fixed for the double mutant is 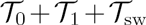. We call this scenario *stochastic tunneling* to distinguish it from other tunneling scenarios that can occur in very large or very heavily subdivided populations. A sufficient condition for stochastic tunneling is that the average time 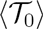 is much larger than both 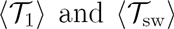. In this case, the total time for the population to adapt is typically dominated by 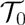. Let *p*_1_ be the probability that a single mutant is successful. Successful single mutants are produced as a Poisson process with rate *N*_T_*µ*_0_*p*_1_, and so the average waiting time for the first successful single mutant is

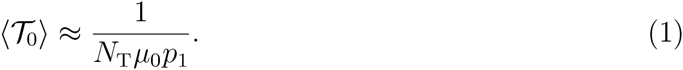

Subdivision accelerates stochastic tunneling by increasing *p*_1_ above its corresponding value in unstructured populations, *p*_1_,_wm_ (the subscript “wm” stands for “well-mixed”). Increases in *p*_1_ are primarily attributable to the effects of competitive assortment, which reduces the rates of selection and drift experienced by single-mutant lineages and so allows them to survive longer and produce more double mutants.

The quantitative effect subdivision has on a given single-mutant lineage depends roughly on the average degree of competitive assortment experienced by the lineage before producing a successful double mutant. We measure competitive assortment with a statistic *F* that quantifies how much more likely a pair of individuals from the same deme are to share a genotype than a pair drawn at random from the total population. The statistic *F* is equivalent to a version of Wright’s fixation index *F*_ST_ defined in Whitlock (2002); it varies between zero and one, with *F* = 0 when a lineage has equal frequency across all demes and *F* = 1 when a lineage is either fixed or absent in each deme. Increasing the degree of subdivision by decreasing the size of demes or the migration rate increases the typical values of *F* experienced by single-mutant lineages and thus increases *p*_1_, Large increases in *p*_1_, of an order of magnitude or more, only occur if successful single-mutant lineages typically fix in one or more demes before producing a successful double mutant and so have average values of *F* close to 1.

Successful single-mutant lineages will only tend to fix within a deme if both *N* and *m* are small. Demes must have substantially fewer than *N*_×_ individuals, where *N*_×_ is the size of an unstructured population in which valley crossing is equally likely to occur by tunneling or sequential fixation. If *N* ≫ *N*_×_, then *p*_1_ ≈ *p*_1_,_wm_, since, even at low migration rates, successful single-mutant lineages remain at low frequency within individual demes and always have small assortments. If *N* ≪ *N*_×_ and *m* is sufficiently small, then successful lineages typically fix within their initial deme and *p*_1_ ≫ *p*_1_,_wm_.

When *N* ≪ *N*_×_, the probability *p*_1_ increases over a wide range of decreasing migration rates, to a maximum 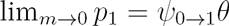, where 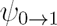 is the probability that a single-mutant lineage fixes in a wild-type deme and *θ* is the probability that a double-mutant lineage that has fixed within a deme goes on to fix in the total population. A sufficient condition for 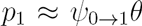 is that migration is rare enough that a single-mutant deme always transitions into a double-mutant deme before being displaced by a wild-type migrant.

Variation in *p*_1_ for higher migration rates depends critically on whether selection against single mutants is strong enough to inhibit their fixing within individual demes. If *δ* ≪ *α/N*, then single mutants are *locally neutral*—they have a neutral probability ≈ 1/*N* of fixing within a deme when migration is rare. (Such mutants are still deleterious with respect to fixation in the total population if *δ* ≫ *α/N*_T_.) If *δ* ≫ *α/N*, then single mutants are *locally deleterious*, and are much less likely than a neutral mutation to fix within a deme due to selection.

If single mutants are locally neutral and the number of demes is large, the effects of subdivision can be found by substituting effective drift and selection parameters into results for unstructured populations. These effective parameters are 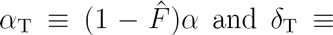 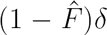, where 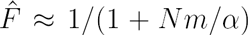 is the equilibrium assortment in the infinite neutral island model. Subdivision has a large effect for small migration rates *m* ≪ *α/N*, for which the effective parameters are greatly reduced below their unstructured values.

If single mutants are locally deleterious, successful single-mutant lineages will either, depending on the migration rate, drift to ~ *α/δ* ≪ *N* copies before producing a successful double mutant, and so never fix within a deme, or drift to ≈ *N* copies and fix within their initial deme. The latter dominate for migration rates *m* ≪ *ηα/N*, where *η* is a constant that is ≪ 1 if *δ* ≫ *α/N*.

Finding the extent to which subdivision accelerates valley crossing requires that we also consider the drift and sweep times. We find that subdivision always increases the average drift and sweep times, by factors at least as large as *p*_1_, Extreme subdivision therefore makes it much more likely for tunneling to be limited by the drift and sweep times. Moreover, these increases in the drift and sweep times can greatly limit the degree to which subdivision accelerates tunneling. The drift time limits how much subdivision can decrease the waiting time for the successful double mutant. We find that subdivision can decrease the average waiting time for a successful double mutant to a minimum of 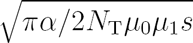. This minimum value is already obtained by an unstructured population if 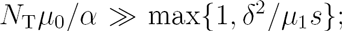 in such cases, subdivision has no effect on the average (or on the distribution) of 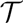. The sweep time is proportional to *α/Nm* for small migration rates *m* ≪ *α/N*, and so will dominate the waiting time for the double mutant to fix in heavily subdivided populations.

Increases in the drift time also limit the degree to which subdivision can increase the probability that the valley is crossed unusually quickly. The probability that a successful double mutant appears by time *t* can only be increased by subdivision to a maximum of 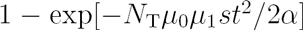. For *t* less than 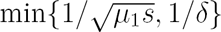, the average drift time in an unstructured population, this maximum occurs even without subdivision and subdivision has no effect on the probability, even if it greatly decreases the average, 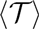.

## 4 Fate of a single-mutant lineage

This section describes how subdivision affects the fate of a single-mutant lineage—in particular, the probability that the lineage is successful and the drift and sweep times. Sections 5 and 6 use these results to find the average and distribution of the waiting time for the population to adapt, including cases where the time is not dominated by the waiting time for a successful single mutant.

### 4.1 Qualitative effects of subdivision

Before presenting our quantitative results for the island model, we first describe the general mechanism by which subdivision increases the probability that a single-mutant lineage is successful as well as the drift and sweep times.

### Subdivision and lineage dynamics

To see how subdivision affects the dynamics of the total copy number of a mutant lineage, we consider a mutant lineage with selection coefficient *f* segregating in an otherwise wild-type population. We let *n*(*t*) be the number of mutant individuals at time *t*, *n*_*i*_ be the number of mutants in the *i*-th deme, and *N*_*i*_ be the size of the *i*-th deme, so that 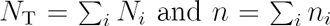. It is often useful to consider frequencies instead of numbers, and so we also let *x* ≡ *n/N*_T_ be the frequency of mutants in the total population and *x*_*i*_ ≡ *n*_*i*_/*N*_*i*_ be the frequency of mutants in the *i*-th deme, so that 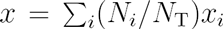. Our goal is to understand how competitive assortment in subdivided populations influences how *n*(*t*) (or *x*(*t*)) changes over time.

We quantify the competitive assortment at time *t* with a measure *F* describing the increase in probability that a pair of competing individuals in the population share a genotype over the probability for a random pair of individuals. First, we consider sampling two individuals with replacement from the total population, and let *J*_*T*_ be the probability that the two individuals share a genotype, or are *identical by state*. Therefore, 
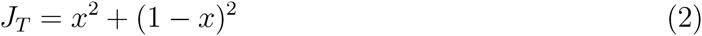
where *x* is the current frequency of the mutation. Next, we consider sampling two individuals by sampling the first from the total population and the second from the set of individuals that compete with the first. For the subdivision models we consider, this set includes all individuals within the deme of the first individual, including the first individual itself. By conditioning on which deme the first individual is chosen from, we can see that

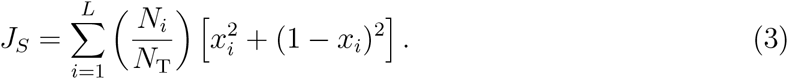

It follows by Jenson’s inequality that if mutants are evenly spatially distributed—that is, if *x*_*i*_ = *x* for all *i*—then *J*_*S*_ = *J*_*T*_, but if mutants are unevenly distributed, then *J*_*S*_ > *J*_*T*_. (This fact is equivalent to the well-known Wahlund effect by which subdivision leads to departures from Hardy-Weinberg proportions in randomly mating diploid species (Charlesworth and Charlesworth 2010).) Intuitively, if mutants are unevenly distributed, they tend to be found in demes with a greater than average frequency of mutants. We define the competitive assortment *F* as the difference between *J*_*S*_ and *J*_*T*_ normalized so that 0 ≤ *F* ≤ 1, or

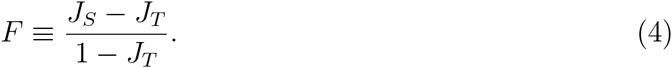

By definition, *F* = 0 when mutants are evenly distributed and *F* = 1 when mutants are either fixed or absent in every deme.

We briefly mention some similarities and differences between the definition (4) and related quantities used in other evolutionary studies. The quantity *F* is equal to *G*_ST_ as defined in Nei (1973) for a biallelic locus if demes are weighted by their size and is also equal to F_ST_ as defined in Whitlock (2002) and Whitlock (2003). We emphasize, however, that our definition differs from other quantities also called F_ST_ and more general “relatedness” or “inbreeding” coefficients (*sensu* Rousset (2002)) in important ways. These quantities as typically defined are deterministic and as such are effectively parameters of the model (Rousset 2002). In contrast, here *F* is a summary statistic of the current state of the population. Some, but not all, of the theoretical knowledge relating to these other quantities carries over when considering the values of *F* that are experienced by mutant lineages. Our reason for the definition of *F* used here is that it provides a direct way of understanding, with minimal assumptions, how competitive assortment affects the evolution of *x*(*t*).

The change in *x* over a short period of time due to selection and drift is characterized by the mean and variance of the change in *x* over one generation. The mean, *M*_*x*_, describes the action of selection, or the deterministic change in *x*, while the variance, *V*_*x*_, describes the action of drift, or the stochastic change in *x*. Given the current state of the population, the mean and variance can be expressed in terms of *x* and *F* (Supplementary Appendix B) as 
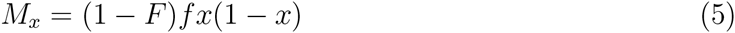
and 
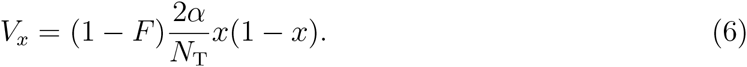

When *F* = 0, Equations (5) and (6) reduce to the familiar expressions for the mean and variance in an unstructured population with mutant frequency *x*. In general, however, both *M*_*x*_ and *V*_*x*_ are reduced by a factor 1 − *F* below their respective unstructured values. This reduction occurs because the fraction of competitive interactions in the population that take place between individuals with different genotypes is reduced by a factor 1 − *F* relative to in an unstructured population.

Even without saying anything about how *F* evolves, we can use Equations (5) and (6) to make some general conclusions about how subdivision affects the long term evolution of the total frequency of the lineage. Starting from a initial frequency *x*(0), whether *x*(*t*) first reaches a frequency *A* before a frequency *B* (with *A* < *x*(0) < *B*) is determined by *M*_*x*_/*V*_*x*_ in the interval between *A* and *B*, and is therefore independent of subdivision (Supplementary Appendix B). This is a more general statement of Muruyama’s well-known result (Maruyama 1970; Maruyama 1974) that the probability that the lineage reaches fixation is independent of subdivision. If we consider the trajectories of *x* that reach *A* from *x*(0) (or *B* from *x*(0)), the frequency will tend to change more slowly in subdivided than in unstructured populations at every intermediate frequency. Compared to an equivalent trajectory in an unstructured population, the time taken to reach *A* (or *B*) will be longer by a factor of 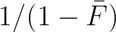, where 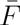 is the average value of *F* over the trajectory (Supplementary Appendix B).

Given our above observations, we can define the probability that a lineage with selection coefficient *f* initially present in *n*_0_ copies reaches *k* copies before going extinct (with 0 ≤ *n*_0_ ≤ *k* ≤ *N*_T_), which we denote *P*_*f*_(*n*_0_,*k*), independently of population structure. This probability can be approximated using standard diffusion methods for unstructured populations (reviewed in Ewens (2004)) and is

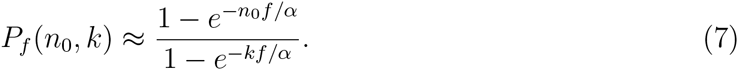

The specific case *n*_0_ = 1 and *k* = *N*_T_ corresponds to the familiar diffusion approximation for the probability of fixation of a mutant lineage when initially present in a single copy.

### Fate of a single-mutant lineage

Subdivision can potentially affect the probability that the single-mutant lineage is successful both through the number of double mutants it produces and through the probability that a double-mutant lineage is successful.

A double-mutant lineage is successful if it is destined for fixation instead of going extinct. The probability that a double-mutant lineage is successful, which we denote *p*_2_, is not entirely independent of subdivision due to the presence of single mutants in the population. If the population were all wild-type, then *p*_2_ = *p*_*s*_(1, *N*_T_) independently of subdivision. For *s* ≫ *α/N*_T_, this probability is ≈ *s/α*, which is usefully interpreted as the reciprocal of the number *α/s* above which the double-mutant lineage is *established*, or very likely to survive drift and sweep through the population (Desai and Fisher 2007). Since single mutants are rare in the total population, in unstructured populations the probability *p*_2_ remains ≈ *s/α*. In subdivided populations, however, single mutants may be frequent in one or more demes and double mutants in primarily single-mutant demes will have a significant extra advantage if *δ* ≳ *s*. In addition, new double-mutant lineages in subdivided populations usually occur in demes with greater than average frequencies of single mutants—if *x* and *F* are the frequency and assortment of single mutants, then the expected frequency of single mutants in the deme where a double mutation occurs is *F* + (1 − *F*)*x*. As a result, subdivision may increase *p*_2_, although we will see that the effect this has on *p*_1_ is usually fairly small.

The number of double mutants produced by a single-mutant lineage is proportional to the total number of single mutant descendants of the lineage, or its *weight* (Weissman *et al*. 2009). The weight *W* is equal to 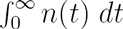, where *n*(*t*) is the number of single mutants at time *t*, and a single-mutant lineage with weight *W* on average produces *Wµ*_1_ double mutants. Subdivision increases the weights of single-mutant lineages, since a lineage is equally likely to reach a maximum of *k* copies in subdivided and unstructured populations, but those in subdivided populations drift for longer in the population. The increase in weight for a given lineage that can be attributable to subdivision grows with the assortment experienced by the lineage, particularly during times when *n*(*t*) is larger.

Overall, subdivision increases the probability *p*_1_ that a single-mutant lineage is successful by increasing *W* without decreasing *p*_2_. The magnitude of the increase in *p*_1_ depends on the assortments experienced by single-mutant lineages; significant increases can only occur if successful lineages tend to have assortments *F* ~ 1. Since *F* is bounded from above the largest frequency of mutants within any individual deme, large assortments require that the lineage has become frequent in at least one deme. Increases in *p*_1_ by an order of magnitude or more require *F* ≈ 1—in other words, successful lineages must typically fix in one or more demes.

Competitive assortment also causes the drift and sweep times to increase in subdivided populations. The drift time increases because single-mutant lineages with high assortments can survive longer and are also less likely to grow very large and produce many double mutants very quickly. The sweep time increases because double-mutant lineages can also have high assortments and so take longer to spread through the population.

## 4.2 Unstructured populations

This section provides the results for unstructured populations needed to describe tunneling in subdivided populations. Previous studies of tunneling in unstructured populations have assumed either the Wright-Fisher model of reproduction, for which *α* = 1/2, or the Moran model of reproduction, for which *α* = 1. Supplementary Appendix C uses an analytical approach similar to Weissman *et al*. (2009) and Weissman *et al*. (2010) to account for general variation in *α*. Here, we summarize the main results and explain the intuition using heuristic arguments also adapted from these earlier studies.

It will be useful later on to distinguish between two ways the drift coefficient, *α*, influences the fate of a single-mutant lineage. First, the drift coefficient determines how quickly the number of single mutants changes by drift. Conditional that a lineage, initially present in a single copy, reaches a number *k* ≲ *α/δ* before going extinct, it will typically do so in ~ *k/α* generations. The lineage will typically then go extinct over an additional ~ *k/α* generations, thereby accumulating a weight ~ *k*^2^/*α* over its lifetime. A single-mutant lineage has a probability ≈ 1/*k* of reaching a number *k* ≪ *α/δ*, but is much less likely to drift to a number *k* ≫ *α/δ;* we can understand this observation by noting that the time required to drift to *k* ≫ *α/δ* is much greater than the time ~ *1/δ* for selection to significantly reduce the lineage number. In order to be able to translate between the time a lineage has survived and the number of the lineage, we also note that a lineage that has survived for *t* ≲ 1/*δ* generations will typically have grown to a number ~ *αt*. These observations follow by extending those of Fisher (2007) and Weissman *et al*. (2009), given for *α* = 1, to general *α*.

The second effect of the drift coefficient occurs in determining the probability *p*_2_ that a double mutant produced by the single-mutant lineage is successful. Recall that the probability *p*_2_ that a double-mutant lineage is successful in an unstructured population is ≈ *s/α*, the reciprocal of the number at which the lineage becomes established. Intuitively, a double-mutant lineage that has drifted to a number ≫ *α/s* is unlikely to drift to extinction within ~ 1/*s* generations, by which time the lineage will have grown substantially from selection. To distinguish the effect of *α* on *p*_2_ from its effect on the rate of stochastic changes in single-mutant lineages, we initially treat the probability *p*_2_ as a new parameter of our model. Later, we substitute *p*_2_ ≈ *s/α* to find results in terms of our original parameters. Meanwhile, our assumption that *µ*_1_ ≪ max{*δ*, *s*} is taken to be equivalent to *µ*_1_ ≪ max{*δ*, *αp*_2_}.

Tunneling dynamics qualitatively differ depending on the strength of selection against single mutants. The probability that a single mutant is successful is

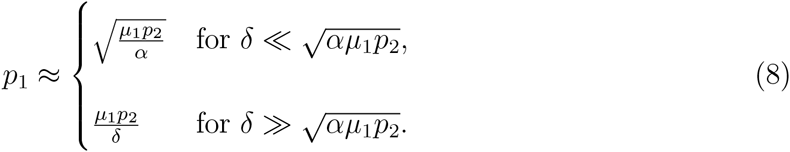

For 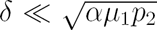, single mutants are (effectively) neutral for tunneling—the probability *p*_1_ does not depend on the strength of selection *δ*. In contrast, if 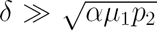, then mutants are deleterious for tunneling and *p*_1_ is inversely proportional to *δ*. The average drift time for neutral and deleterious single mutants is

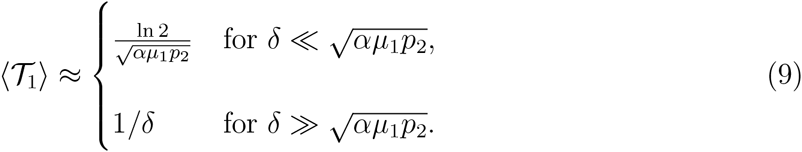

The full distribution of 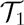 is found in Supplementary Appendix C. The critical feature of this distribution is that a successful lineage is very unlikely to drift for much fewer or much more than 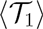 generations. Instead, successful lineages typically drift for ~ 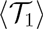 generations and reach a number ~ *α*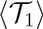, before producing a successful double mutant.

We can understand the expressions for *p*_1_ and 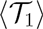 in (8) and (9) in terms of the behavior of typical successful lineages when single mutants are either neutral or deleterious. Conditional on a single-mutant lineage having weight *W*, it produces at least one successful double mutant with probability 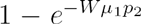. Lineages typically become successful in one of two ways (Weissman *et al*. 2009; Weissman *et al*. 2010): either they have a large weight *W* ≳ 1/*µ*_1_*p*_2_ and thus are nearly assured to produce at least one successful double mutant, or they have a small weight *W* ≪ 1/*µ*_1_*p*_2_, yet manage to produce exactly one successful double mutant. Which strategy is used depends on whether or not selection is strong enough to limit the single-mutant lineages to small numbers and thus to small weights.

Successful neutral lineages follow the first strategy of having a large weight: they drift to numbers 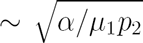 over 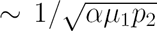 generations, reaching a weight ~ 1/*µ*_1_*p*_2_. The probability *p*_1_ in (8) corresponds to the probability of drifting to such a number, while 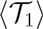 in (9) roughly corresponds to the time required to do so. These lineages strike a sweet spot for producing a successful double mutant; lineages that only reach smaller numbers have much smaller weights, while lineages that reach larger numbers have larger weights but are not significantly more likely to produce a successful double mutant. The condition 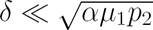 ensures that the lineages can drift to a number 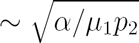 neutrally (equivalently, it ensures that the time 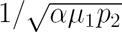 required to do so is smaller than the time ~ 1/*δ* for selection to purge the lineage (Weissman *et al*. 2009)).

If 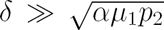, selection limits single-mutant lineages to much smaller numbers and weights, forcing successful deleterious lineages to follow the second strategy. Since almost all deleterious lineages have weights ≪ 1/*µ*_1_*p*_2_, we can approximate the probability that a lineage produces a successful double mutant as 〈*W*〉*µ*_1_*p*_2_, where 〈*W*〉 is the average weight of a deleterious lineage and is approximately 1/*δ*. The dominant contribution to 〈*W*〉, and thus to *p*_1_, comes from lineages that drift to numbers ~ *α/δ* over periods of ~ 1/*δ* generations. Although lineages that drift to larger numbers tend to have larger weights, these lineages are so unlikely that they make a negligible contribution *p*_1_. These observations explain the expressions *p*_1_ ≈ *µ*_1_*p*_2_/*δ* in (8) and 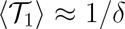 in (9).

Substituting *p*_2_ = *s/α* into (8) and (9) gives expressions for *p*_1_ and 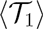 in terms of our basic model parameters. These expressions are 
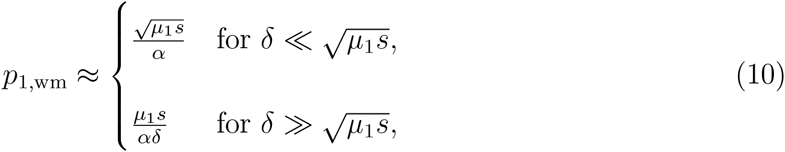
and 
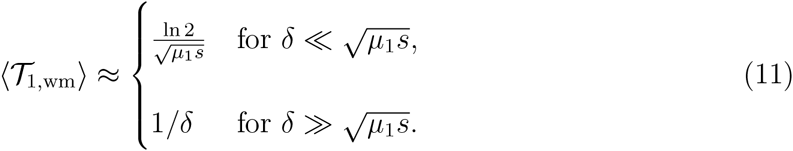

We have used the notation *p*_1,wm_ in (10) and 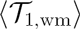 in (11) to enable comparison between these expressions and our results for *p*_1_ and 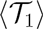 in subdivided populations (the subscript “wm” stands for “well-mixed”).

We conclude our discussion of tunneling in unstructured populations by returning to our assumption that *µ*_1_ ≪ max{*δ*, *s*} (or that *µ*_1_ ≪ max{*δ*, *αp*_2_}). Equations (10) and (11) show that this assumption is equivalent both to *p*_1_ ≪ *s/α* and to 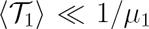. If instead *µ*_1_ ≫ max{*δ*, *s*}, then *p*_1_ ≈ *s/α* and 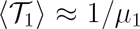; in other words, *p*_2_ ≈ *s*/*α* forms an upper bound for *p*_1_ and 1/*µ*_1_ an upper bound for 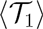. By assumption, these upper bounds are not met if the population is unstructured, but we will see that they can be reached if the population is sufficiently subdivided.

### Sequential fixation in small populations

In sufficiently small populations, successful lineages are likely to fix in the population before producing a successful double mutant and valley crossing will occur by sequential fixation. More generally, we can compare the probabilities that a single mutant is successful by the tunneling and sequential pathways to determine which is more likely. Since we do not allow back mutation, a single-mutant lineage that reaches fixation always produces a successful double mutant. Therefore, the probability that a single-mutant lineage is successful by sequential fixation is simply the probability that the lineage drifts to fixation, equal to *P*_−*δ*_(1, *N*_T_) or approximately

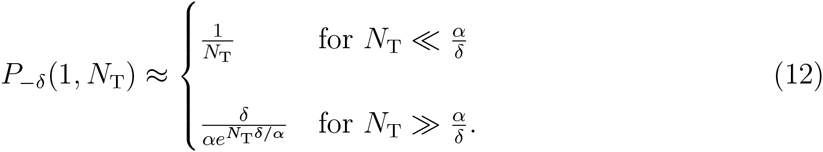

After previous authors (Weinreich and Chao 2005; Weissman *et al*. 2009), we define the threshold population size *N*_×_ as the size where the probability *p*_1_ of success by tunneling equals *P*_−*δ*_(1, *N*_T_). If *N*_T_ ≫ *N*_×_, then *p*_1_ ≫ *P*_–*δ*_(1, *N*_T_) and tunneling is much more likely than sequential fixation, while the opposite is true if *N*_T_ ≪ *N*_×_. The threshold size if single mutants are neutral or deleterious for tunneling, found by assuming the approximation (10) for *p*_1_, is

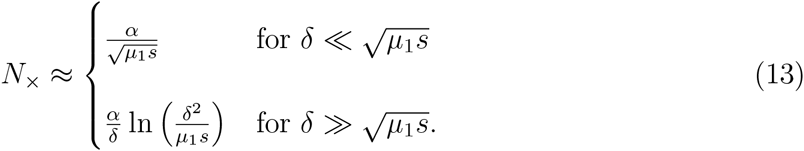

Equation (13) is valid if *N*_×_ ≫ *α/s*, so that *p*_2_ ≈ *s*/*α* for *N*_T_ ~ *N*_×_. This requirement is always met if *δ* ≲ *s*, but in general may not be; see Supplementary Appendix D for the general solution for *N*_×_. For neutral single mutants, *N*_×_ equals the typical copy number 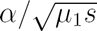 reached by successful lineages in the tunneling pathway. For deleterious single mutants, *N*_×_ is smaller than 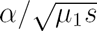, but is still substantially larger than the typical number *α/δ* reached by successful tunneling lineages. When *N*_T_ ≈ *N*_×_ in the deleterious case, the fact that selection greatly limits the probability that a lineage drifts to *N*_T_ copies is exactly offset by the assurance of producing a successful double mutant if it does so.

## 4.3 Limit as *m* → 0 and the isolated-demes regime

Demes must be small enough for subdivision to increase the probability that a single mutant is successful. Otherwise, successful single-mutant lineages will produce a successful double mutant without ever reaching high frequency in a deme, even under extremely restricted migration. The only significant effect of subdivision in such cases is possibly increasing the sweep time.

Here, we determine how small demes must be for subdivision to potentially increase *p*_1_ by a large amount by considering the fate of single-mutant lineages in the limit *m* → 0—the best case scenario for subdivision to increase *p*_1_. Valley-crossing dynamics are easily studied in the limit, since a single-mutant lineage must either go extinct or produce a double mutant that fixes in the initial deme before migration in or out of that deme can occur. Once fixed within the deme, a double-mutant lineage has a probability *P*_*s*_(*N*, *N*_T_) ≡ *θ* of then fixing in the total population, approximately equal to

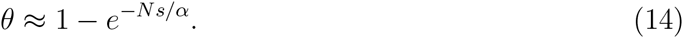

As in unstructured populations, we consider two ways a single-mutant lineage might produce a successful double mutant—by tunneling within the deme, or by first reaching fixation in the deme. The probability that a single mutant is successful by tunneling within the deme is approximately *p*_1_,_wm_, the probability of tunneling in an unstructured population of size *N*_T_, while the probability the mutant is successful by first fixing in the deme is *P*_−*δ*_(1, *N*)*θ*. The two are approximately equal when *N* = *N*_×_, and for demes with much more than *N*_×_ individuals, tunneling is much more likely, while for demes with much fewer than *N*_×_ individuals, fixation is much more likely.

At least a fraction of demes must have sizes *N* ≪ *N*_×_ for subdivision to have a large effect on single-mutant lineages at low migration rates. Since we are interested in determining when subdivision can significantly accelerate valley crossing, we assume *N* ≪ *N*_×_ for the remainder of our analysis. We will also assume that *N*_T_ ≫ *N*_×_, so that tunneling is the dominant mode of valley crossing at high migration rates. This assumption simplifies our analysis by ensuring that single-mutant lineages remain rare in the total population across all migration rates; we describe these lineages as tunneling with respect to the total population even if they fix in one or more demes. It is also possible for subdivision to accelerate valley crossing when *N*_T_ ≪ *N*_×_. Although we do not explicitly consider this situation, it can be analyzed straightforwardly by combining our results for the rate of tunneling in subdivided populations to that of sequential fixation, which is unaffected by subdivision.

Over the next two sections, we will determine how *p*_1_ and 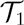 vary as a function of migration rate, *m*, while the size, *N*, of demes is fixed so that *N* ≪ *N*_×_ ≪ *N*_T_, and find it useful to keep in mind the above dynamics as occurring for sufficiently low migration rates. These dynamics are characterized by new single-mutant lineages having a probability 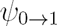 of fixing within their initial deme and, once fixed, being guaranteed to produce a double-mutant deme before being replaced by a wild-type migrant or colonizing additional demes. Since migration does not play a role in determining the fate of these lineages, we refer to the range of parameters where such dynamics apply the *isolated-demes regime*.

In the isolated-demes regime, the probability that a single mutant is successful is approximately 
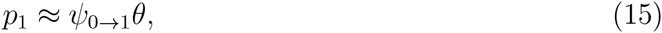
and the drift time is approximately exponentially distributed with mean 
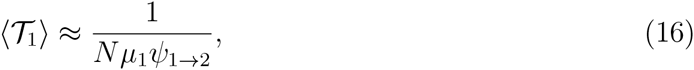
which follows since the single-mutant deme produces double mutants at rate *Nµ*_1_ that each have a probability 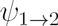 of fixing in the deme. These expressions for *p*_1_ and 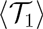 are decreasing in *N* and are significantly greater than *p*_1_,_wm_ and 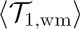 if and only if *N* ≪ *N*_×_. Both (15) and (16) become insensitive to *N* at small numbers—in particular, *p*_1_ ≈ *s/α* and 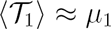 for *N* ≪ min{*α/δ*, *α/s*}. These values for *p*_1_ and 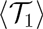 are the maximum that occur as the degree of subdivision is increased by decreasing *N* and *m*.

We conclude our discussion of the isolated-demes regime by determining the range of migration rates for which it applies. To do so, we use the fact that when *m* and *N* are sufficiently small, we can approximate fixation or loss of a lineage that has entered a deme by migration or mutation as being practically instantaneously. Therefore, we approximate a single-mutant lineage that has fixed in its initial deme by a *deme birth-death (DBD) process*, in which single-mutant demes can give birth by colonizing a new deme, die by being replaced by a wild-type migrant, and mutate into a double-mutant deme. To simplify notation for the rates of these events, we let 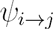 denote the probability that an individual with *j* mutations fixes within a deme currently occupied by individuals with *i* mutations. For example, a single-mutant lineage fixes in a wild-type deme with probability 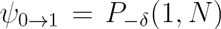 and a wild-type lineage fixes in a single-mutant deme with probability 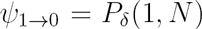. So long as single mutants are rare in the total population, each single-mutant deme gives birth at rate 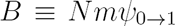, dies at rate 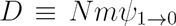, and mutates at rate 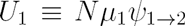 into a double-mutant deme. The isolated-demes regime corresponds to migration rates where *U*_1_ ≫ max{*B*, *D*}, ensuring that a single-mutant deme always mutates before dying or giving birth; since *D* ≥ *B*, this condition is equivalent to *U*_1_ ≪ *D*. We define *m*\* as the migration rate where *U*_1_ = *D*, given by

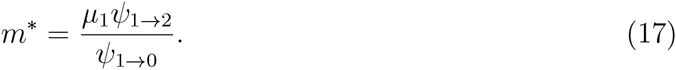

The isolated-demes regime corresponds to *N* ≪ *N*_×_ and *m* ≪ *m*\*.

## 4.4 Locally neutral single mutants (*δ* ≪ *α/N*)

How single-mutant lineages are impacted by low migration rates critically depends on whether or not these lineages can fix neutrally within a deme, making valley-crossing dynamics most easily understood by considering locally neutral and locally deleterious single mutants separately. This section presents results for *p*_1_ and 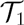 when single mutants are locally neutral and the next presents results when single mutants are locally deleterious. The intermediate scenario that occurs when *δ* ~ *α/N* is analyzed in Supplementary Appendix F.

### Summary of methods

Analyzing valley crossing by modeling the frequency dynamics within each deme is intractable; therefore, we must develop approaches for approximating lineage dynamics. We develop two primary methods for predicting valley crossing when single mutants are locally neutral by extending previous methods for approximating frequency dynamics in the island model. The 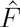 method, outlined below and detailed in Supplementary Appendix E, predicts *p*_1_ and 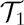 when single-mutant lineages are locally neutral and typically drift to ≫ *N* copies before producing a successful double mutant. The DBD method, described in Section 4.3 and Supplementary Appendix C, predicts *p*_1_ and 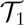 when single-mutant lineages typically fix in their initial deme before producing a successful double mutant. For 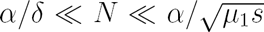, the two methods are valid for distinct but overlapping ranges of migration rates, together predicting dynamics across the full range of *m*.

Before describing how these methods are used, we make some useful observations about successful single-mutant lineages. By assumption, the deme size *N* is smaller than the number 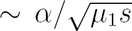 reached by successful neutral lineages and the number ~ *α*/*δ* reached by successful deleterious lineages in unstructured populations. Successful single-mutant lineages therefore drift to numbers ≫ *N* before producing a successful double mutant at high migration rates. They only drift to ≈ *N* copies at migration rates *m* ≪ *m*\*; however, lineages that reach fewer than *N* copies always have a negligible probability of being successful. Consequently, so long as *m* ≪ *α/N*, we can assume that successful single-mutant lineages fix in their initial deme. In addition, we expect a wide range of migration rates *m*\* ≪ *m* ≪ *α/N* over which successful lineages fix in their initial deme, but drift to ≫ *N* copies, fixing in multiple demes, before producing a successful double mutant. (Note that this situation contrasts with that for locally deleterious single mutants, which drift to numbers ~ *α/δ* ≪ *N* at high migration rates.)

The 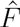 method reduces the effects of subdivision to a constant reduction in the rates of selection and drift experienced by single-mutant lineages. In particular, we model the single-mutant lineage in the subdivided population by one in an unstructured population of equal total size, but with effective parameters 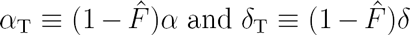, where 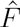 is a constant that approximates the long-run average assortment of locally neutrally lineages. Meanwhile, since the establishment probability of the double mutant is approximately unaffected by subdivision for *δ* ≪ *α/N*, we keep *p*_2_ = *s/α*. In this way, predictions for *p*_1_ and 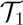 follow directly from the theory for unstructured populations described in Section 4.2 and Supplementary Appendix C.

Over sufficiently long periods, the average assortment of locally neutral lineages tends towards a value that is independent of both the lineage frequency in the total population and the strength of selection against the lineage. This value is approximately equal to the equilibrium assortment at a polymorphic neutral locus with no mutation in the infinite island model, consisting of an infinite number of demes of *N* individuals. In the neutral infinite island model, frequency at the locus remains constant in the population as a whole; however, the frequency within each deme changes due to drift and migration, causing the assortment to deterministically reach an equilibrium value, 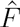. The equilibrium value equals the probability that a pair of alleles sampled from the same deme, looking backward in time, coalesce to a common ancestor before either migrates to a different deme. Intuitively, a pair that coalesce must be identical, while a pair that migrates resembles a sample from the total population at a short time in the past, but where the population has the same total frequency as the present; thus, 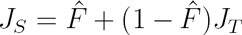. The equilibrium 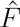 is approximated by Wright’s well-known formula (Wright 1951) 
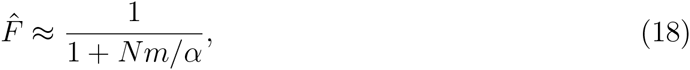
which follows immediately from the observation a pair of lineages coalesce within a deme at rate 2*α/N* and migrate at rate 2*m*. Thus, large assortments of *F* ~ 1 in the neutral, infinite island model correspond to migration rates *m* ≲ *α/N*.

Unlike in the infinite model, the assortment in the finite model is stochastic; however, if *L* ≫ 1, *δ* ≪ *α/N*, and the lineage is large enough, then the average assortment of the lineage approximates 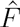. Recall that *F* is bounded by maximum local frequency of the lineage; thus, mutant lineages with numbers 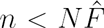 necessarily have assortments smaller than 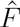. However, the assortments of locally neutral lineages with numbers 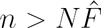 tend to fluctuate around a value 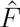, such that averaged over a increasing number of generations the assortment becomes closer and closer to 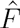 (Figure 3 and Supplementary Appendix E). Weak local selection does not affect the assortment of these lineages with 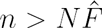 because selection takes 1/*δ* generations to affect genealogies within demes, while assortment dynamics are governed by the faster processes of coalescence and migration within demes (Roze and Rousset 2003).

**Figure 3:**
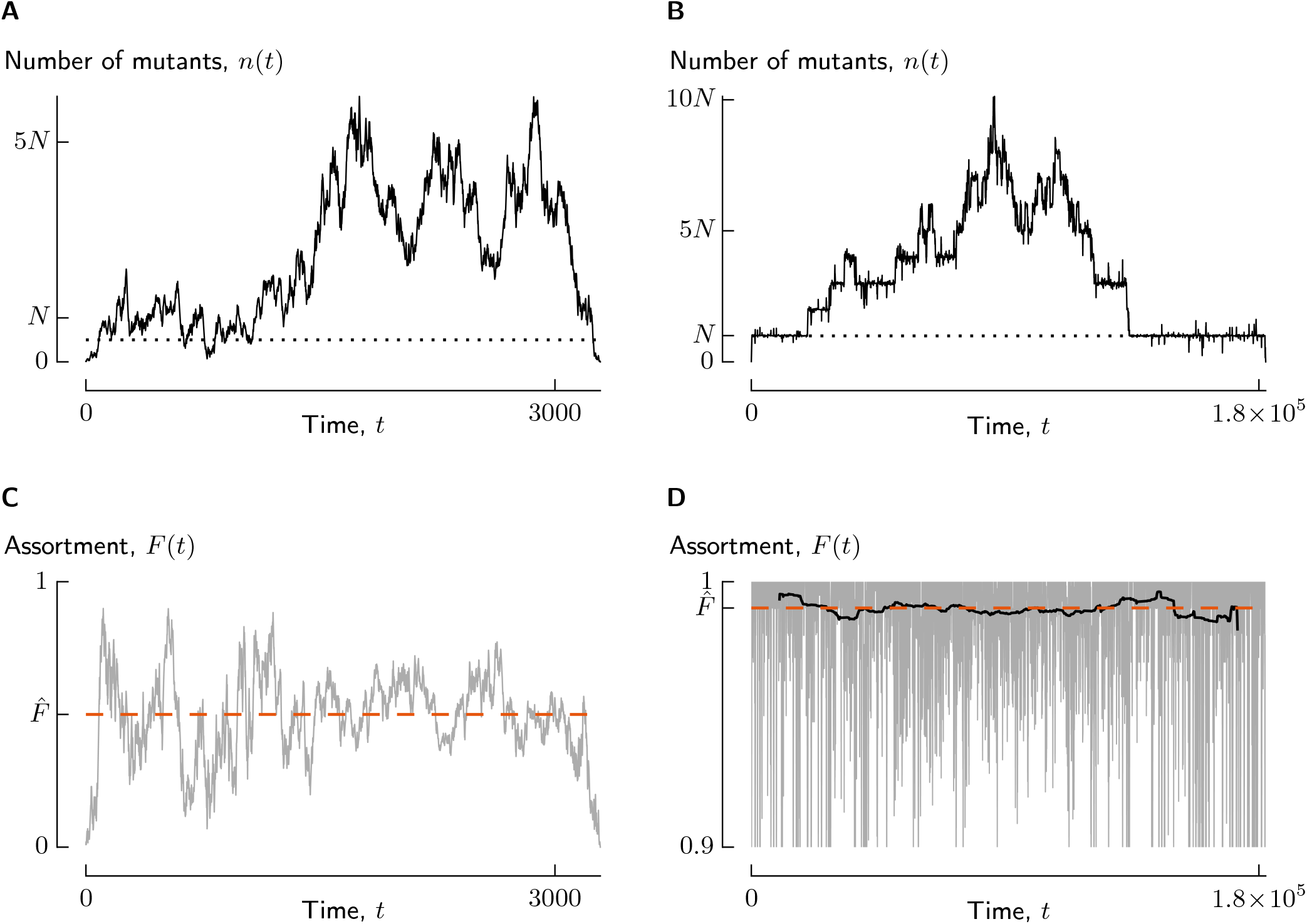
The assortment of a locally neutral lineage fluctuates around 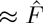 while the lineage is present in more than 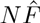 copies. Shown are the trajectories of *n* and *F* for a locally neutral lineage (*Nδ/α* = 0.2) at each of two migration rates, corresponding to *Nm/α* = 1 (Panels A and C) and *Nm/α* = 10^−2^ (Panels B and D). Each lineage began as a single copy and reached at least *α/δ* = *5N* copies before going extinct. In Panels A and B, the solid line shows *n*(*t*) while the dotted line marks the line 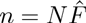. In Panels C and D, the grey line shows *F* and the dashed orange line indicates 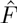, while the black line in Panel D shows the centered moving average of *F* over a window of 1/m generations. Note that assortments below 0.9 are not shown in Panel D. Parameters are *L* = 100, *N* = 100, *α* = 0.5, and *δ* = 10^−3^, with *m* = 5 × 10^−3^ for A and C and *m* = 5 × 10^−5^ for Band D.

On the basis of similar observations, previous authors suggested approximating the dynamics of frequency in the total population by taking 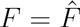 (Cherry and Wakeley 2003; Roze and Rousset 2003; Wakeley 2003; Whitlock 2003; Wakeley and Takahashi 2004); however, conditions for when the stochastic fluctuations in *F* can be ignored for predicting lineage dynamics were not completely described. Several authors (Cherry and Wakeley 2003; Roze and Rousset 2003; Wakeley 2003) argued the constant *F* approximation is valid when *L* is sufficiently large that fluctuations in *F* are negligible. However, small fluctuations in *F* can require that the lineage is spread over an extremely large number of demes. Lineages that begin as a single copy necessarily occupy a relatively small number of demes and have significant variation in *F* even for large *L* (e.g., Figure 3 and Supplementary Figure S4). In Supplementary Appendix *E*, we find that a constant *F* approximation accurately predicts lineage dynamics if *n* ≫ *N*, but may not for smaller *n*. The condition *n* ≫ *N* ensure that even large fluctuations in *F* typically average out over the time periods required for significant changes in *n* by selection or drift (Supplementary Appendix E). A single-mutant lineage that drifts to a maximum number *k* ≫ *N* typically spends a majority of its life-time (and accumulates an even larger majority of its weight) at numbers ≫ *N*. Thus, if successful lineages typically drift to ≫ *N* copies, we can use the constant *F* approximation to model tunneling dynamics.

A second method for approximating dynamics at low migration rates is suggested by the fact that for *m* ≪ *α/N*, fixation or loss of a migrant within a demes occurs approximately independently of further migration. If successful single-mutant lineages are likely to reach at least *N* copies and fix in their initial deme, then we can approximate lineage dynamics by treating fixation and loss within a deme as instantaneous and model the number of single-mutant demes by the DBD process described in Section 4.3. We assume that lineages that reach fewer than *N* copies make a negligible contribution to *p*_1_, a valid assumption since 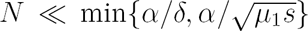. Predictions for valley-crossing dynamics under the DBD approximation are derived in Supplementary Appendix C. These predictions agree with those for the isolated-demes regime for *m* < *m*\*. They also agree with predictions from the 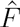 method at migration rates *m*\* ≪ *m* ≪ *α*/*N*—at these migration rates, successful lineages always fix in their initial deme, but tend to reach numbers ≫ *N*, fixing in multiple demes, before producing a successful double mutant. Consequently, we use the 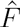 method and isolated-demes approximation primarily for understanding the main quantitative and qualitative effects of subdivision, and use the DBD method results mainly for verification and for combining with the 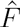 method to make numerical predictions across full range of migration rates. Simulations show that numerical predictions for *p*_1_ and 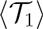 found by these methods are very accurate across all migration rates for *Nδ/α* as large as 0.2 (Figure 5 and Supplementary Figure S1).

### Results

The relationships of *p*_1_ and 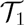 with *m* depend on whether double mutants are locally beneficial or neutral (*s* ≫ *α/N* or *s* ≪ *α/N*), and we first present results when they are locally beneficial. When *δ* ≪ *α/N* and *s* ≫ *α/N*, tunneling dynamics fall into one of five distinct parameter regimes. These regimes can be conveniently visualized in a phase diagram as a function of migration rate *m* and the single mutant selection coefficient *δ* (Figure 4). The five regimes occupy the region *δ < α/N* in Figure 4; the two additional regimes with *δ > α/N* are described in the next section. The probability that a single mutant is successful in these five regimes is approximately 
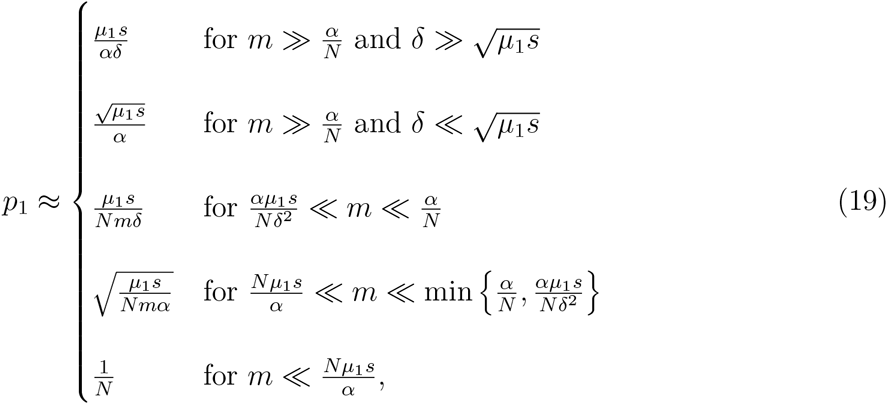
and the average drift time is approximately 
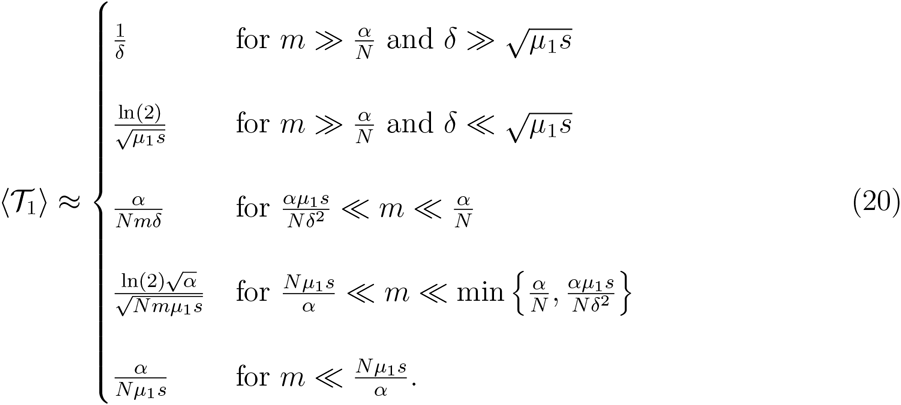

**Figure 4:**
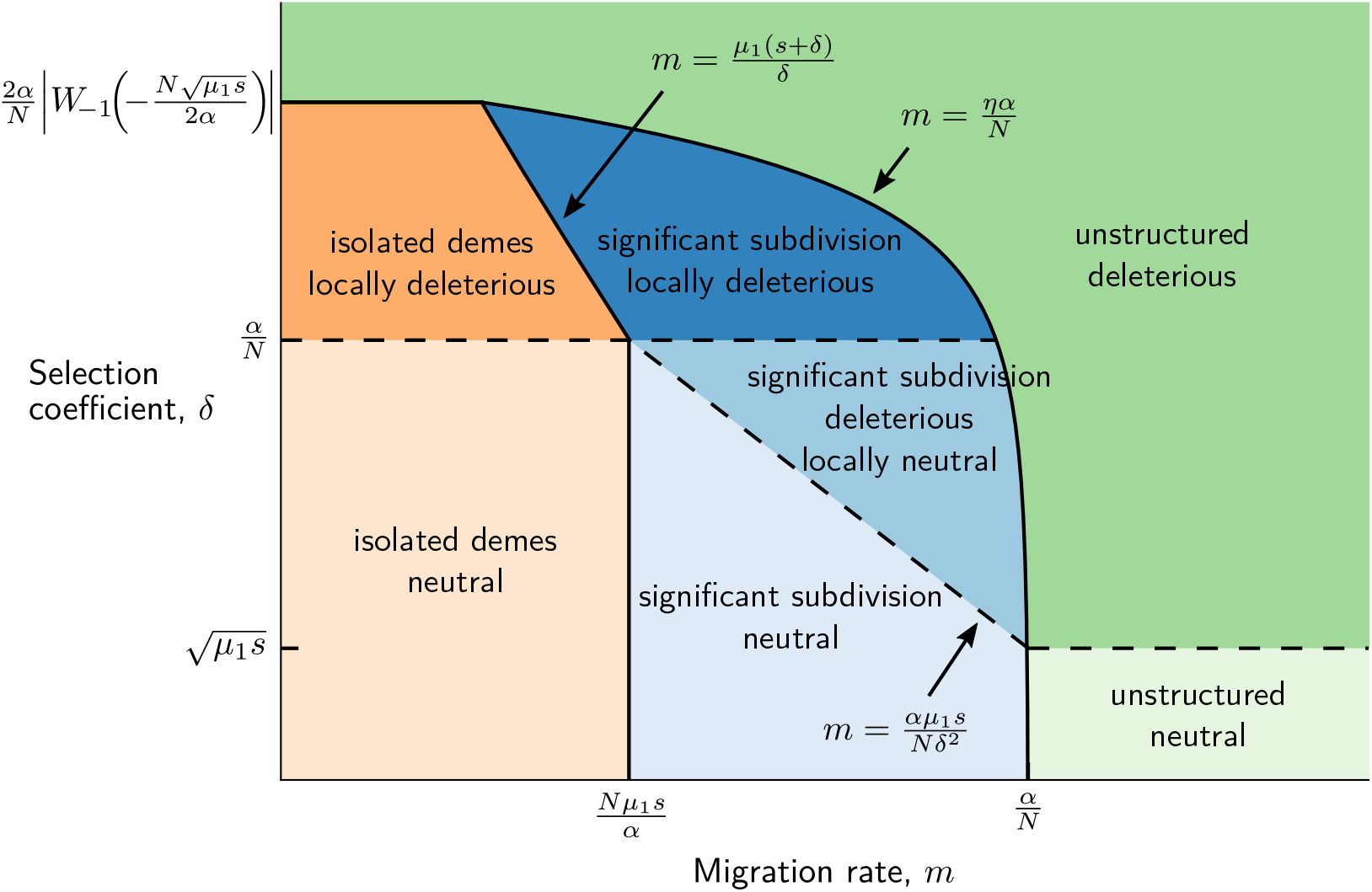
The dynamics of successful single mutants separate into seven regimes depending on the migration rate, *m* and the selection coefficient of the single mutant, *δ*, when demes are intermediately sized with *α/s* ≪ *N* ≪ 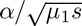, so that effects of subdivision are possible and double mutants always spread after fixing within a deme. Both axes are on a logarithmic scale, and *W*_−1_ is the negative real branch of the product-log function. To make the figure, we fixed *N* = 10^3^, *α* = 0.5, *µ*_1_ = 10^−8^, and *s* = 0.05 and varied *m* and *δ*.

Complete analytical predictions are given in Supplementary Appendix C. Figure 5 shows our predictions for *p*_1_ and 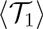 as a function of *m*, along with estimates from simulations, in a case where 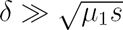, so that single mutants are deleterious for tunneling at high migration rates, and Supplementary Figure S1 shows a case where 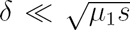. In these figures, the range of migration rates corresponding to each regime is color coded according to Figure 4.

**Figure 5:**
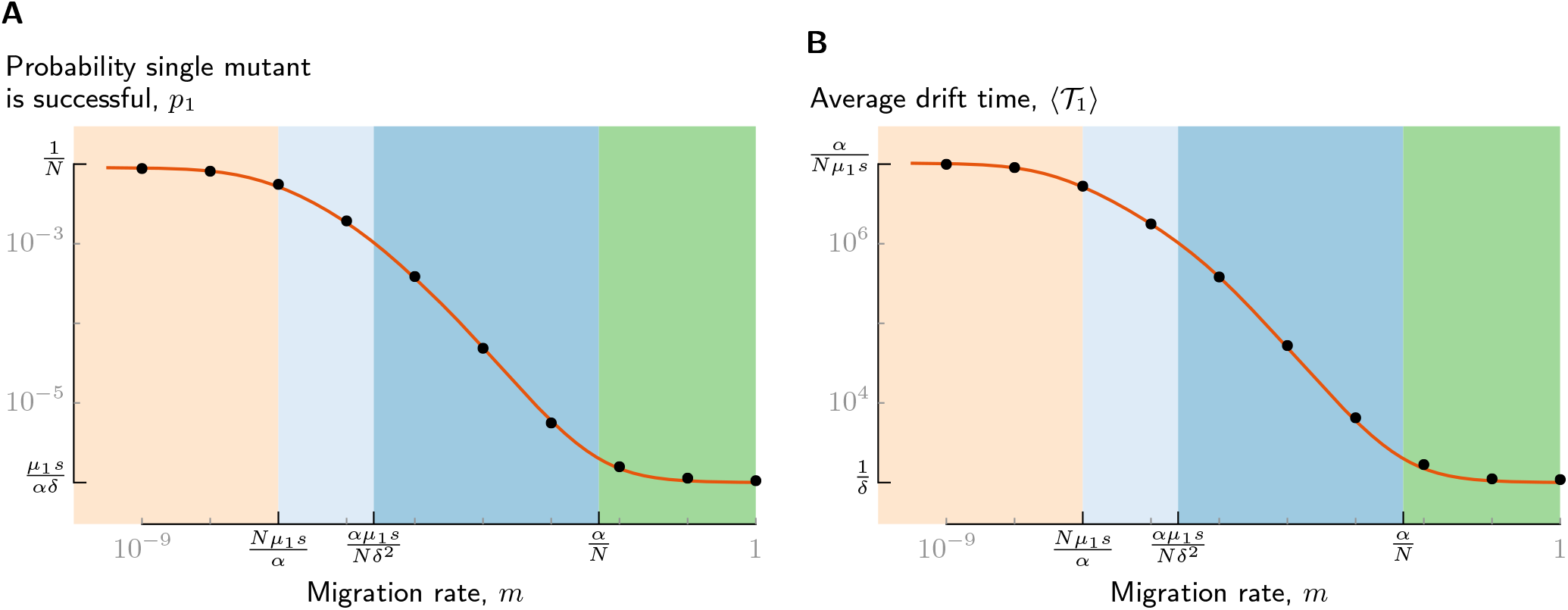
When single mutants are locally neutral (*δ* ≪ *α/N*), our analytical predictions using the 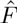 and DBD methods accurately predict similar increases in the probability that a single mutant is successful, *p*_1_, and the average drift time, 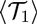, as the migration rate, *m*, decreases. Here, single mutants are deleterious for tunneling at high migration rates 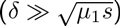 and double mutants are locally beneficial (*s* ≫ *α/N*), so that there are four regimes for *p*_1_ and 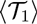 depending on *m*. In each panel, the orange line shows the predictions for *p*_1_ and 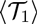 by smoothly joining the 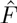 and DBD approximations. The dots show estimates from simulations (95% confidence intervals are smaller than the dots). Background colors correspond to the regimes labeled in Figure 4. Both axes have a log scale, with orders of magnitude delineated by small tick marks. Large tick marks indicate the regime boundaries on the *x*-axis and the limiting values of *p*_1_ and (71) on the *y*-axis as given in Equations (19) and (20). Parameters are *L* = 100, *N* = 100, *α* = 0.5, *µ*_1_ = 10^−8^, *δ* = 10^−3^, and *s* = 0.05, so that *Nδ/α* = 0.2, *Ns/α* = 10, and *δ*^2^/*µ*_1_*s* = 2×10^3^.

These results show that *p*_1_ and 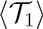 show little variation with *m* at very high and very low migration rates, and increase significantly with decreasing *m* over a range of intermediate migration rates. For *m* ≫ *α/N*, migration is sufficiently frequent that the population is effectively unstructured for tunneling. The condition *m* ≫ *α/N* corresponds to 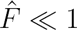; since single-mutant lineages have small assortments, subdivision has almost no effect on tunneling dynamics. The two regimes with *m* ≫ *α/N* in (19) and (20) simply correspond to whether single mutants are deleterious or neutral for tunneling in an unstructured population. Low migration rates *m* ≪ *Nµ*_1_*s/α* correspond to the isolated-demes regime described in Section 4.3. For *α/δ* ≪ *N* ≪ *α/s*, the various fixation probabilities described in Section 4.3 are approximately 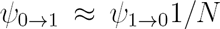 and *θ* ≈ 1, so that *m*\* ≈ *Nµ*_1_*s/α* and *p*_1_ ≈ 1/*N* and 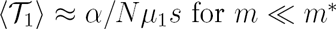.

Over the range of intermediate migration rates *α/Nµ*_1_*s* ≪ *m* ≪ *α/N*, both *p*_1_ and 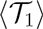 increase with a decreasing migration rate from their unstructured values to their isolated-demes values. The mathematical relationship of *p*_1_ and 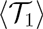 on *m* is easily understood using the 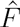 method, substituting the effective drift and selection coefficients *α*_T_ and *δ*_T_ into Equation (8) for *p*_1_ and (9) for 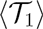. Noting that 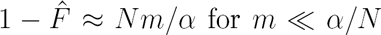, we see that *α*_T_ and *δ*_T_ are each reduced by a factor *Nm/α* as compared to their unstructured values. Accordingly, for single mutants that are deleterious for tunneling, *p*_1_ and 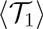 are increased by a factor *α/Nm* relative to their unstructured values, while for single mutants that are neutral for tunneling, *p*_1_ and 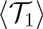 are increased by a factor of 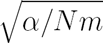. Whether single mutants are neutral or deleterious for tunneling is determined by whether *δ*_T_ is smaller or larger than 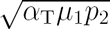. Single mutants with 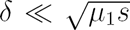 are neutral across all migration rates. Single mutants with 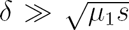 are deleterious for 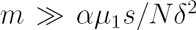 and neutral for *m* ≪ *αµ*_1_*s/Nδ*^2^; intuitively, for migration rates below *αµ*_1_*s/Nδ*^2^, subdivision has reduced the number a lineage must reach to have a weight ~ 1/*µ*_1_*p*_2_ to less than *α/δ*. Figure 5 shows *p*_1_ and 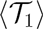 in this second case where 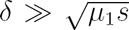, so that, as *m* decreases, single mutants transition from being deleterious to neutral for tunneling.

When *s* ≪ *α/N*, we must modify the above results as follows. Equations (19) and (20) continue to predict *p*_1_ and 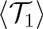 for migration rates *m* ≫ *µ*_1_ min{*α/Nδ*, *α/Ns*}, but not for lower migration rates. For rates *m* ≪ *µ*_1_ min{*α/Nδ*, *α/Ns*}, we instead find that *p*_1_ ≈ *s/α* and 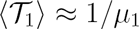; thus, both *p*_1_ and 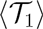 are approximately independent of further decreases in *m*. The isolated-demes regime corresponds to *m* ≪ *µ*_1_. Therefore, when *s* ≪ *α/N, p*_1_ and 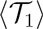 reach their maximum values at migration rates outside of the isolated-demes regime and where single-mutant lineages still reach ≫ *N* copies before becoming successful. This observation contrasts with what we observed when *s* ≫ *α/N*, where *p*_1_ and 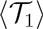 increased with decreasing *m* until *m*\*. Intuitively, this difference is because for *s* ≪ *α/N*, both *p*_l_ and 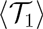 reach their theoretical upper bounds of *p*_2_ and 1/*µ*_1_, respectively, once *m* ≪ *µ*_1_ min{*α/Nδ, α/Ns*}, and so further decreases in *α*_T_ and *δ*_T_ have no effect.

As Figure 5 makes apparent, when single mutants are locally neutral, subdivision increases *p*_1_ and 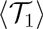 by similar proportions. Our analytical results show that if *δ* ≪ *α/N*, then 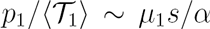 across all regimes, and the same relationship between *p*_1_ and 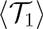 applies in unstructured populations. Therefore, increasing the degree of subdivision by decreasing either *N* or *m* has a near-constant trade-off between decreasing 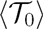 and increasing 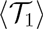 once *N* is smaller than *α/δ*. We consider the consequences of this trade-off for the mean and distribution of the valley-crossing time in Sections 5 and 6.

## 4.5 Locally deleterious single mutants *(δ ≫ α/N)*

When single mutants are locally deleterious, at migration rates *m* ≪ *α/N*, valley crossing may still be dominated by successful lineages that never reach high frequency within a deme. Our assumption that 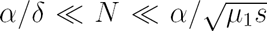 implies 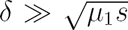, so that single mutants are deleterious for tunneling in unstructured populations. Therefore, successful lineages drift to ~ *α/δ* copies at high migration rates, not nearly enough to fix within a deme in the absence of migration. And yet, our assumption that *N* ≪ *N*_×_ ensures that successful lineages reach *N* copies and fix within a deme for sufficiently low migration rates. For lineages to fix within their deme (and for subdivision to increase *p*_1_), the benefit of fixing in a deme-potentially gaining a much larger weight than lineages that reach only ~ *α/δ* copies—must outweigh the cost of a low probability of drifting to *N* copies.

### Summary of methods

To solve valley crossing dynamics when single mutants are locally deleterious, we separately consider the contributions of lineages that drift to numbers ≲ *α/δ* and those that drift to at least *N* copies. Lineages that reach intermediate numbers *k* such that *α/δ* ≪ *k* ≪ *N* do not contribute to valley crossing for the same reason as in large unstructured populations—compared to lineages reaching ~ *α/δ* copies, their lower probability of occurrence greatly outweighs their slightly higher weights. Only lineages that reach *N* or more copies and fix within a deme have a sufficient weight advantage to potentially offset their low probability. For brevity, we refer to single-mutant lineages that reach a maximum number ≲ *α/δ* as *type A* and those that reach *N* or more copies as *type B*. Note that type-B lineages only have a possibility of influencing valley crossing at migration rates low enough that they will fix in their initial deme. We define 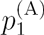 as the contribution to *p*_1_ from type-A lineages; i.e., the probability that a new lineage is type A, multiplied by the probability that a type-A lineage is successful, and similarly define 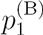. The probability that a single-mutant is successful follows from the separate contributions as 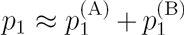. In addition, we let 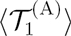 be the average drift time of successful type As and 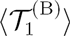 be that of type Bs. The overall average drift time 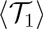 is given by the average of 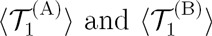 weighted by 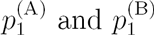, respectively.

Type-A lineages have small assortments and so act similarly to deleterious lineages in unstructured populations. Successful type-A lineages have a maximum assortment of ~ *α/Nδ*, set by the maximum frequency a lineage present in ~ *α/δ* copies can obtain in any one deme. At migration rates *m* ≪ *δ*, we expect successful lineages to remain within their initial deme and thus have *F* ~ *α/Nδ*. This small positive assortment leads to an increase in *p*_1_ on the order of *α/Nδ* percent (Supplementary Appendix F analytically confirms this prediction). To a good approximation, we can ignore this small effect of structure in determining the fate of a type-A lineage. Since the vast majority of single-mutant lineages are type A, the approximate contribution to *p*_1_ and 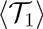 from type As are simply given by *p*_1_ and 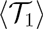 for deleterious tunneling in an unstructured population; i.e., 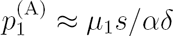 and 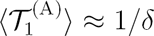.

Since significant contributions from type-B lineages are limited to low migration rates ensuring fixation in the initial deme, we approximate their dynamics using the DBD model described in Section 4.3 and Supplementary Appendix C. The contribution to *p*_1_ from type Bs equals the probability 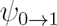 that a lineage fixes in its deme, times the probability that a single-mutant deme is successful via the DBD process, producing a successful double-mutant deme before going extinct. The dynamics of the DBD process simplify, however, for *δ* ≫ *α/N*, since the deme birth rate is much smaller than the deme death rate *D*, allowing us to ignore the possibility that the lineage spreads beyond its initial deme before being driven extinct by an immigrating wild-type lineage (Supplementary Appendix C). The probability that a single-mutant deme is successful is thus approximately *U*_1_/(*D* + *U*_1_), the probability that the deme mutates before going extinct, times the probability *θ* that the double-mutant deme is successful. The contribution to *p*_1_ from type-B mutants is therefore

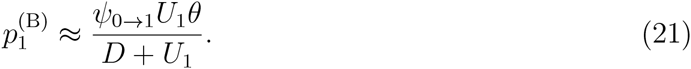

The drift time of successful type-B lineages is approximately exponentially distributed with average

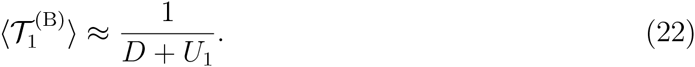

We recall that the migration rate *m*\* determining the boundary of the isolated-demes regime is given by migration rate where *D* = *U*_1_, For migration rates *m* ≪ *m*\*, where *D* ≪ *U*_1_, the expressions (21) and (22) for 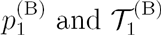, respectively, reduce to the values of *p*_1_ and 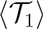 in the isolated-demes regime (i.e., 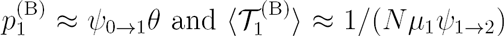. For migration rates *m* ≫ *m*\*, for which *D* ≫ *U*_1_), these expressions can be approximately written as 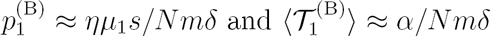, where 
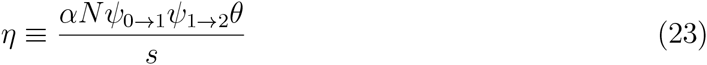
is a constant that is ≪ 1 for *δ* ≫ *α/N* and ≈ 1 for *δ* ≪ *α/N*.

### Results

Combining these predictions for the two types of lineages completely describes the fate of single-mutant lineages across the full range of migration rates. The relative size of *s* and *α/N* determines the maximum values of *p*_1_ and 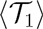 as *m* decreases by determining *θ*, but, unlike when *δ ≪ α/δ*, does not qualitatively change the relationships of *p*_1_ and 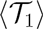 with *m*. Figure 6 shows predictions for *p*_1_ and 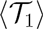 and the contribution from the two types in a case where *Nδ/α* = 8 (large enough for a clear separation type-A and B lineages), and *Ns/α* = 10 (large enough that *θ* ≈ 1). As expected, *p*_1_ and 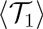 are driven by type-A lineages at high migration rates and by type-B lineages at low migration rates. The two types contribute equally to *p*_1_ (i.e., 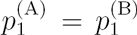) at a migration rate *m ≈ ηα/N*, so that for 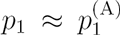 at migration rates *m ≫ ηα/N* and 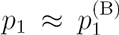 at rates *m* ≪ *ηα/N*. The migration rate must therefore be significantly lower for subdivision to increase *p*_1_ than when *δ* ≪ *α/N*. Due to the much longer drift times of type Bs at rates *m ≪ α/N*, type Bs dominate 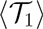 over a wider range, 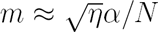, than they do *p*_1_. In other words, for rates 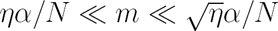, the average drift time approximates 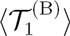 and is much larger than 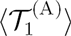, even though the majority of successful mutants are type A.

**Figure 6:**
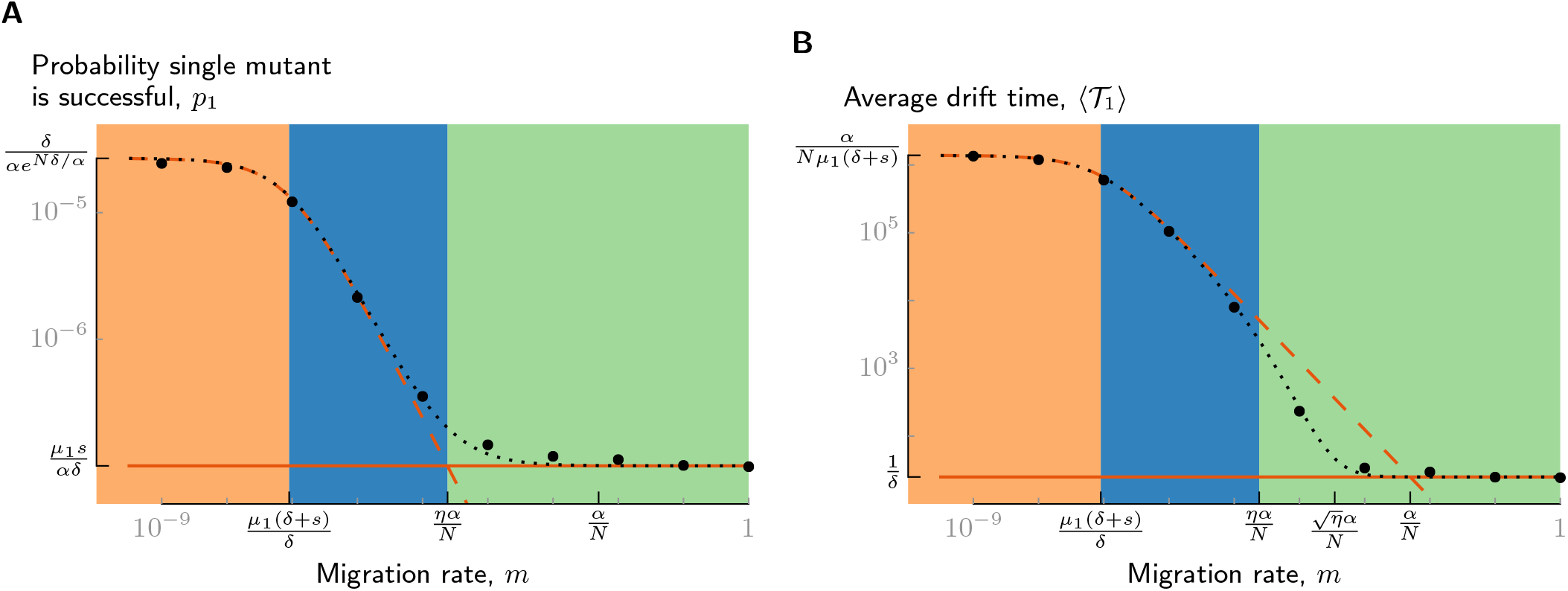
When single mutants are locally deleterious (*δ* ≫ *α/N*), increases in the probability that a single mutant is successful, *p*_1_, and the average drift time, 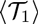, at low migration rates are due to the contributions of type-B mutants. Solid orange lines show predictions for type-A lineages 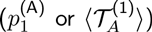 dashed orange lines show predictions for type-B lineages 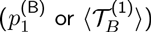 and the dotted black line shows predictions for *p*_1_ and 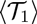 accounting for both types. Large black dots show estimates from simulations (95% confidence intervals are smaller than the dots). Background colors correspond to the regimes labeled in Figure 4. Both axes have a log scale, with orders of magnitude delineated by small tick marks. Large tick marks indicate the regime boundaries on the *x*-axis and the limiting values of *p*_1_ and 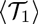 on the *y*-axis as given in Equations (24) and (25). Parameters are *L* = 50, *N* = 100, *α* = 0.5, *µ*_1_ = 4 × l0^−8^, *δ* = 0.04, and *s* = 0.05, so that *Nδ/α* = 8 and *Ns/α* = 10.

There are three distinct regimes for the probability that a single mutant is successful, mapped in the region *δ > α/N* in Figure 4 and colored in Figure 6. In these regimes, the probability that a single mutant is successful is approximately

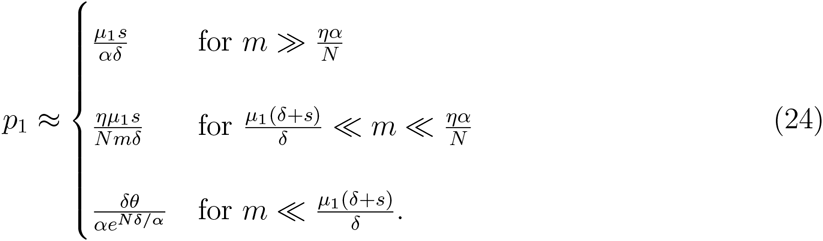

The regime *m* ≫ *ηα/N* corresponds to migration rates where type-A mutants make the dominant contribution to *p*_1_ and so tunneling dynamics are similar to deleterious tunneling in an unstructured population. In the other two regimes, type Bs dominate *p*_1_ such that 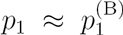 and is significantly increased over its unstructured value. The regime *m* ≪ *µ*_1_(*δ* + *s*)/*δ* corresponds to the isolated-demes regime; here, the fraction 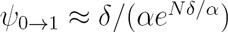 of mutants that are type B are successful with probability *θ*. In the intermediate regime *µ*_1_(*δ* + *s*)/*δ* ≪ *m* ≪ *ηα/N*, type-B mutants dominate but only have a probability *U*_1_/*D* of producing a double-mutant deme, causing *p*_1_ ∝ 1/*m*. The drift time has four regimes-the three regimes for *p*_1_ plus the additional regime 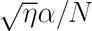 described above—in which the average drift time is approximately

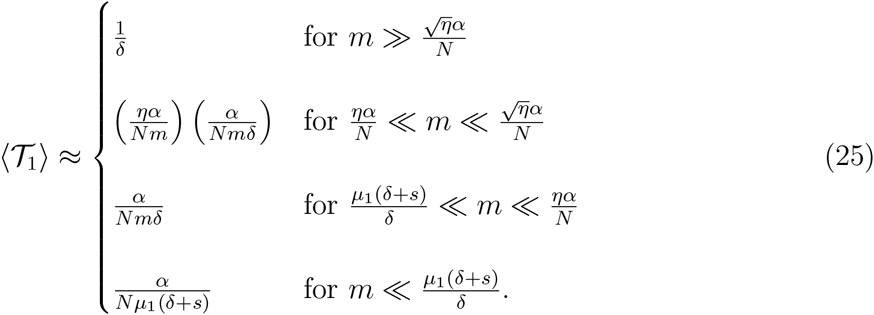

Subdivision tends to increase 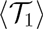 much more than *p*_1_ at migration rates low enough for subdivision to have a significant effect. The larger increase can be traced to the fact that 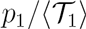 equals *µ*_1_*s/α* for type-A lineages but equals *ηµ*_1_*s/α* for type-B lineages. Therefore, an even steeper trade-off between 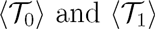 occurs for locally deleterious single mutants than for locally neutral.

We conclude this section with an observation about the distribution of drift times when single mutants are locally deleterious. The distribution of 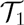 has two humps, centered at 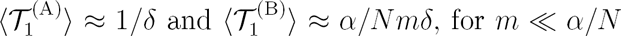. These humps can be seen most clearly when *m ≈ ηα/N*, so that successful lineages are equally likely to be of either type (Figure 7). For *m* ≪ *ηα/N*, the hump from successful type As forms a tiny fraction of the overall distribution. But, if valley crossing is limited by the drift time, the shorter drift times of type-A lineages may allow them to dominate valley crossing. We return to this point when considering the mean and distribution of 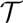 in the next two sections.

**Figure 7:**
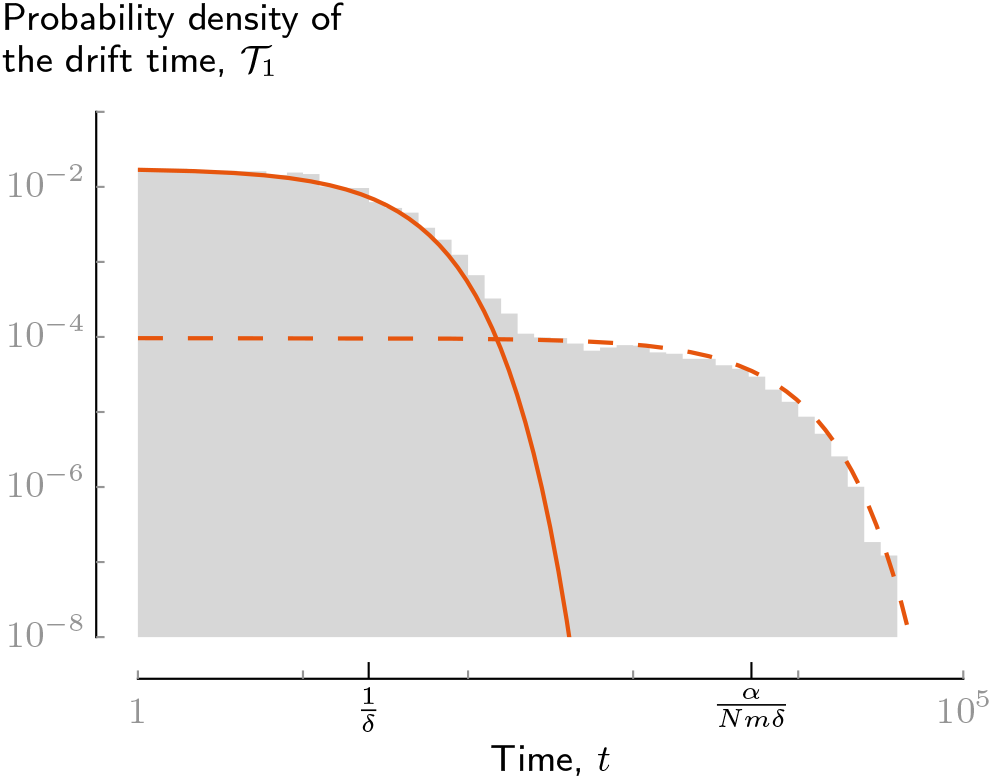
When single mutants are locally deleterious (*δ* ≫ *α/N*), the distinct behavior of type-A and type-B successful single mutants can be seen in the two distinct humps in the distribution of drift times when *m* ~ *ηα/N*. The hump at times ~ *1/δ* is from successful type-A mutants, while the hump at ~ *α/Nmδ* is from successful type-B mutants. The solid and dashed orange lines show predictions for the distributions for type-A and type-B mutants, respectively, while the grey area is a histogram of simulation results. Parameters are as in Figure 6, but with *m* = 2.4×10^−5^ ≪ *ηα/N*, so that a successful mutant is equally likely to be type A or type B.

## 4.6 Sweep time

After a single-mutant lineage produces a successful double mutant, it takes an additional 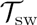 generations for the double-mutant lineage to become fixed in the population. Supplementary Appendix G reviews sweep dynamics in unstructured populations and develops new approximations for sweeps in the island model. In unstructured populations, the average sweep time is approximately

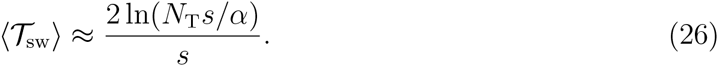

The dominant factor in this expression for 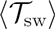 is 1/*s*, the timescale over which selection significantly increases the number of double mutants. In island populations at migration rates *m* ≪ *α/N*, the average sweep time is approximately

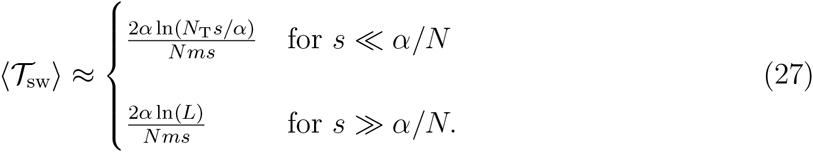

In (27), the dominant factor is *α/Nms*, which is sets the timescale for selection to significantly increase the number of double mutants when *m* ≪ *α/N*. See Supplementary Appendix G for more detailed expressions and interpretations further showing how subdivision influences the sweep. Equation (27) shows that heavy subdivision can lead to large sweep times even in modestly sized populations. Comparing (27) to (26) and ignoring logarithmic terms shows that subdivision increases 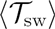 by a factor 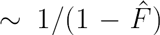, which is always similar to or larger than the factor of increase in *p*_1_. In Section 5, we consider how the long sweep times in heavily subdivided populations can limit the ability for large amounts of subdivision to accelerate adaptation.

## 5 Average waiting time for the population to adapt

This section uses the results from Section 4 to describe the average waiting time 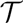 for a successful double mutant to arise and start to spread and the average waiting time for complete fixation of the double mutant. If these waiting times are typically dominated by 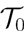, then increases in *p*_1_ due to subdivision directly translate into decreases in the average time for the population to adapt. However, in very large or heavily subdivided populations, the drift and sweep times can form a substantial fraction of the adaptation time. Since subdivision increases the drift and sweep times, in such cases the decreases in the time to adapt will be substantially less than the increase inp, We first present our results for 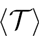 for subdivided populations with *δ* ≪ *α/N* (together with unstructured populations), then for subdivided populations with *δ* ≫ *α/N*, and finally consider the overall time for the double mutant to fix, accounting for the sweep.

### Unstructured populations and subdivided populations with *δ* ≪ *α/N*

The average waiting time for a successful double mutant in unstructured populations and in subdivided populations for which single mutants are locally neutral are both approximately given by (Supplementary Appendix C)

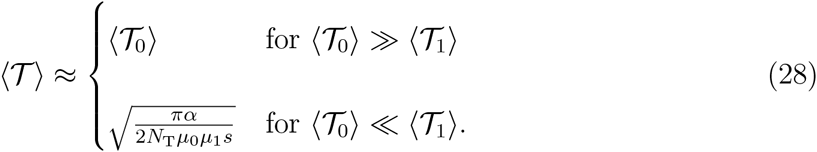

The effects of subdivision in (28) are encompassed solely by the determination of 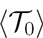 and 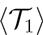.

If 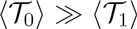, then the first successful single-mutant lineage that occurs in the population will typically produce a successful double mutant before another successful single mutant can arise. In this case, we will typically have that 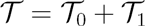 and so 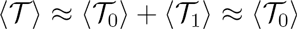. If 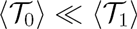, then the approximation 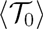 significantly underestimates 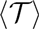 by ignoring the drift time. But, since multiple successful single-mutant lineages are likely to appear before the first successful double mutant, the approximation 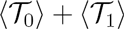 significantly overestimates 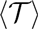—if many successful single-mutant lineages segregate simultaneously, the lineage that is first to produce a successful double mutant must do so much more quickly than usual.

Weissman *et al*. (2009) called the regime in unstructured populations with 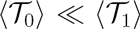 the *neutral semi-deterministic tunneling (NSD)* regime. In the NSD regime, the total number of single mutants in the population grows as ≈ *N*_T_*µ*_0_*t* until the time 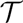 when a successful double mutant is produced. Our results from Section 4.2 show that the condition 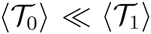 corresponds to *N*_T_*µ*_0_/*α* ≫ max{1, *δ*^2^/*µ*_1_*s*} in unstructured populations. The condition N_T_*µ*_0_/*α* ≫ 1 ensures that the number of single mutants only shows small fluctuations about its expectation, while *N*_T_*µ*_0_/*α* ≫ *δ*^2^/*µ*_1_*s* ensures that a successful double mutant is produced within 1/*δ* generations, before negative selection has an effect. In the NSD regime, the role of drift only affects the probability *p*_2_ ≈ *s/α* that double mutants establish. From the fact that successful double mutants are produced at rate ≈ *N*_T_*µ*_0_*µ*_1_*st/α* it is easily shown that the average waiting time for a successful double mutant is 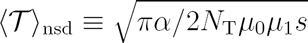.

In Supplementary Appendix C, we show if *δ* ≪ *α/N* and 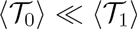, then NSD dynamics also occur in subdivided populations and 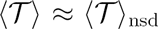. By decreasing 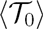 and increasing 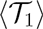, subdivision greatly expands the situations where NSD dynamics occur. This finding makes intuitive sense given our discussion in Section 4.1 of how subdivision reduces the rates of drift and selection acting on the number of single mutants in the population.

Since the approximation for 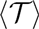 in (28) is roughly given by the maximum of 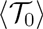 and 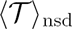, the time 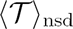 sets a lower bound on 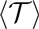 regardless of the degree of subdivision in the population. One can imagine increasing the degree of subdivision by decreasing *N* and/or *m* while keeping the total population size fixed, causing 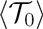 to decrease and 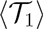 to increase. If the population is small enough that *N*_T_*µ*_0_/*α* ≪ *µ*_1_/*s*, eventually *p*_1_ will reach its theoretical maximum of ≈ *s/α* and 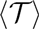 will reach ≈ *α/N*_T_*µ*_0_*s* while the drift time is still negligible. However, if *N*_T_*µ*_0_/*α* ≫ *µ*_1_/*s*, then 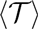 can only decrease to a minimum of 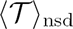 This minimum roughly occurs at the point where 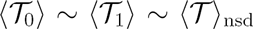, which can be seen as minimizing 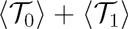 subject to the constraint 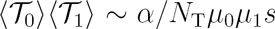 that arises from the trade-off between *p*_1_ and 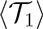. Figure 8 shows 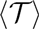 in this scenario as the migration rate is decreased and *N* is held fixed. The average 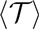 reaches 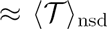 and stops decreasing at a migration rate that is still much larger than the migration rate *m*\* at which *p*_1_ stops increasing.

**Figure 8:**
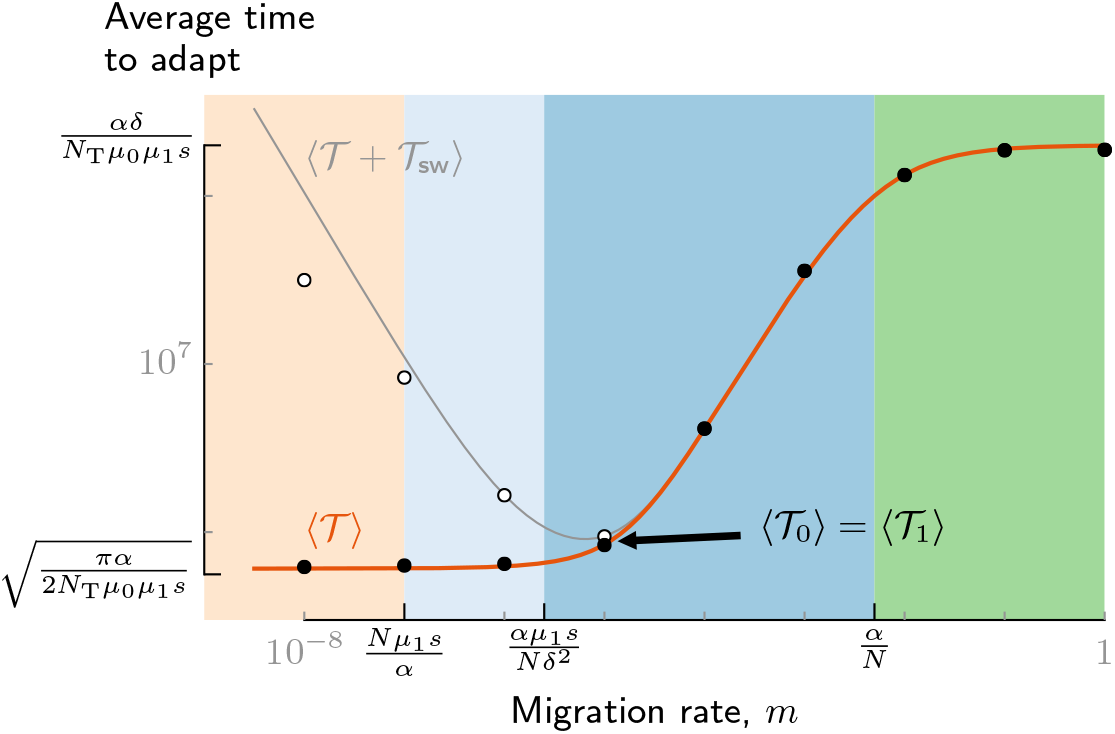
The drift and sweep times limit how much extreme subdivision decreases the average time for the population to adapt. Shown is the average waiting time for a successful double mutant, 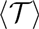, (prediction: orange line; simulation results: black dots) and the average waiting time for the double mutant to fix in the total population (prediction: grey line; simulation results: white dots). Single mutants are locally neutral (*Nδ/α* = 0.2); parameters, *x*-axis, and background colors are the same as in Figure 5, while *µ*_0_ = 5 × 10^−7^, so that *N*_T_*µ*_0_/*α* = 10^−2^. For these parameters, the migration rate where 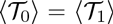 is coincidentally near that where 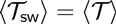.

We conclude our discussion of 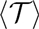 when *δ* ≪ *α/N* by observing that if *N*_T_*µ*_0_/*α* ≫ max{1, *δ*^2^/*µ*_1_*s*}, then NSD dynamics already occur in the absence of population structure, and no degree of subdivision can effect a decrease in 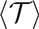.

### Subdivided populations with *δ* ≫ *α/N*

When single mutants are locally deleterious, we find 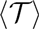 by separately considering the average waiting times for a successful double mutant to first be produced by a type-A or a type-B lineage. If 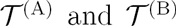 are the waiting times for a successful double mutant to be produced by a type-A and by a type-B lineage, respectively, then 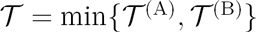. The averages and distributions of 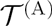 and 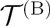 are derived in Supplementary Appendix C. For most parameter choices, one of 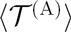 or 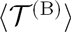 will be significantly smaller than the other, and the type with the smaller average almost always produces the first successful double mutant. In this case, the average waiting time 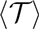 is approximately given by the minimum of 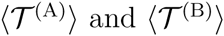.

If the total waiting time is dominated by the waiting time for the first successful single mutant, then the condition for 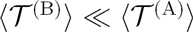 is simply that *m* ≪ *ηα/N*. But, due to the much longer drift times of type Bs, this condition is in general not sufficient. We let 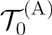 be the waiting time for the first successful type-A single mutant and 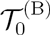 be the waiting time for the first successful type-B single mutant. Both 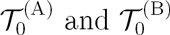 are exponentially distributed with rates 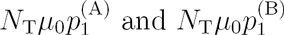, respectively, so that successful type Bs occur at a faster rate when *m < ηα/N*. Since type-A lineages act like deleterious lineages in an unstructured population, the average waiting time for a successful double mutant from a type-A lineage is roughly given by the maximum of 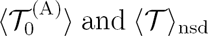 (see Equation (28)). The average waiting time for a successful double mutant from a type-B lineage is approximately (Supplementary Appendix C)

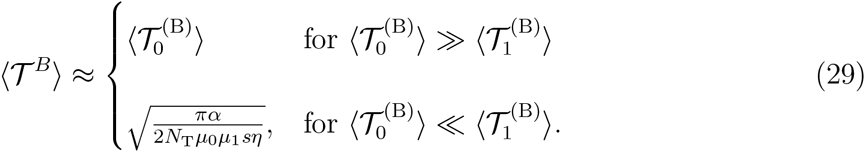

The average waiting time in (29) is approximately given by the maximum of 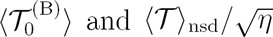

Comparing these results for 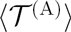 and 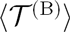 shows that 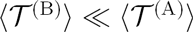 if and only if 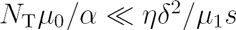 and *m* ≪ *ηα/N*. Since 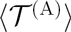 approximately equals 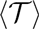 in an unstructured population, these conditions are also the conditions for subdivision to significantly decrease 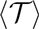 when *δ* ≫ *α/N*. The requirement on *N*_T_*µ*_0_/*α* can be understood by noting that if N_T_*µ*_0_/*α* ≫ *ηα/N*, then 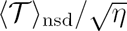 is much larger than 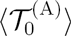. In this case, tunneling by type-B lineages remains slower than by type-As even at migration rates *m* ≪ *ηα/N* due to the much longer drift times of type-B lineages.

### Conditions for a significant reduction in 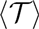

These results show that for subdivision to significantly decrease the average waiting time for a successful double mutant, not only must demes and migration rates be small enough, but the scaled rate of new single mutants entering the population, *N*_T_*µ*_0_/*α*, must not be too large. Explicit conditions for a significant decrease in 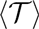 across all regimes for the island model are that *N* ≪ *N*_×_ and *m* ≪ *ηα/N*, which ensure that *p*_1_ ≫ *p*_1,wm_, and that *N*_T_*µ*_0_/*α* ≪ min{1,*ηδ*^2^/*µ*_1_*s*}, which ensures that longer drift times do not completely offset larger values of *p*_1_ in the subdivided population.

### Total time to fixation

For approximating the total time until complete fixation of the double mutant, there are two scenarios to consider. The first scenario is that all or a vast majority of the final population traces its ancestry back to the first successful double mutant; in this scenario, the sweep is said to be *hard* (Messer and Petrov 2013). If a hard sweep is likely, the average waiting time until complete fixation of the double mutant is approximately 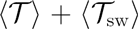. The second scenario is that multiple successful double-mutant lineages arise and make significant contributions to the final population; in this scenario, the sweep is said to be *soft* (Messer and Petrov 2013). If a soft sweep is likely, the approximation 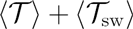 may substantially overestimate the total time to fixation since the contributions from later successful double mutants tend to shorten the time between the arrival of the first successful double mutant and fixation. A necessary (but insufficient) condition for soft sweeps is that 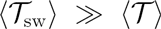 (Supplementary Appendix G). Supplementary Appendix G provides more details about the fixation process under both the hard and soft scenarios.

By decreasing 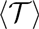 and increasing 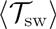, subdivision makes the sweep time more likely to limit the total time to fixation of the double mutant. The fastest rate of adaptation typically occurs at intermediate levels of subdivision where 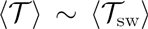. An example is shown in Figure 8, where both the average time to fixation is shown as a function of migration rate in addition to 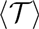. The time to fixation begins increasing with decreasing *m* below the point where 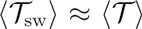 (coincidentally, this happens to be near the migration rate where 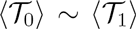. This figure also illustrates the overestimation of the time to fixation by the hard-sweep approximation at low migration rates.

## 6 Distribution of the waiting time

In biological applications such as the development of cancer or drug resistance, rare instances where a population adapts atypically quickly may be of utmost importance. For these applications, it is critical to consider not only the average but also the distribution of 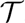, or the probability 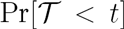 that a successful double mutant has occurred by a specific time *t*. Supplementary Appendix C finds the probability distribution of 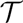 for unstructured populations and subdivided populations with locally neutral or locally deleterious single mutants. This section summarizes these results and uses them to find the conditions in which subdivision significantly increases 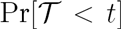 as well as the maximum values of 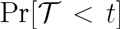 that can be obtained by subdividing a population.

### Unstructured populations and subdivided populations with *δ* ≪ *α/N*

As with the mean, the distribution of 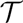 can be described similarly for unstructured populations and subdivided populations when single mutants are locally neutral. When 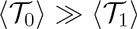, so that 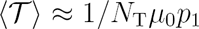, the probability that a successful double mutant has occurred by *t* is approximately

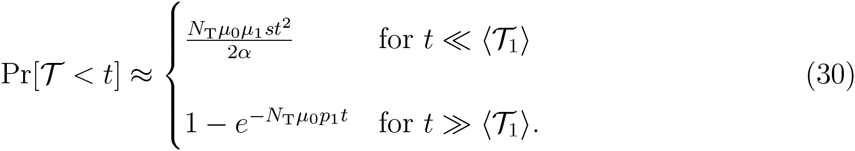

At times 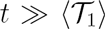, the distribution of 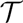 is approximately exponential with mean 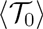, as at these times, 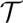 is dominated by the waiting time for the first successful single mutant (i.e., the drift time is negligible). However, the probability that a successful double mutant appears by a time 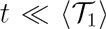 is much less than under the exponential approximation, since tunneling at these times requires a single-mutant lineage that is extra lucky—not must the lineage produce a successful double mutant, it must do so much more quickly than a typical successful lineage.

The expression for the probability in (30) at times 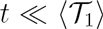 can be understood using a simple argument. Let *n*_1_(*t*) be the total number of single mutants in the population at time *t* and 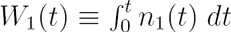 be the total weight from all single-mutant lineages up to *t*. Early on, the total weight from single mutants tends to be small. If *t* is small enough that we can safely assume that *W*_1_(*t*) ≪ 1/*µ*_1_*p*_2_, then 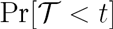 is approximately 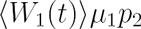. Initially, the average number of single mutants grows simply as 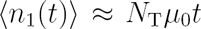, independently of population structure. Thus, early on, the average total weight grows as 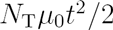 and the probability of at least one successful double mutant as 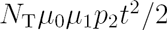. This argument fails for 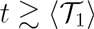, at which point enough time has passed for selection and mutation to double mutants have reduced the growth of 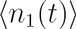 or for *W*_1_ to become ≳ 1/*µ*_1_*p*_2_ (Supplementary Appendix C). Weissman *et al*. (2010) gives an alternate informal derivation for the approximation for 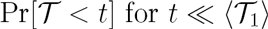 in unstructured populations based on the likely dynamics of lineages that become successful within *t* generations.

Equation (30) shows that even if subdivision greatly reduces 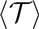 relative to an unstructured population, it only significantly increases 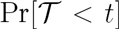 for times much longer than the average drift time in the unstructured population (Figure 9). The range of times in which 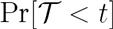 is unaffected by subdivision is substantial if single mutants are close to neutral (as in the figure) and so have long drift times even when the population is effectively unstructured. This effect can be seen as another consequence of the trade-off between *p*_1_ and the drift time. Subdivision increases *p*_1_ by causing lineages to survive longer than they otherwise would, but also causes these lineages to grow more slowly in number and weight.

**Figure 9:**
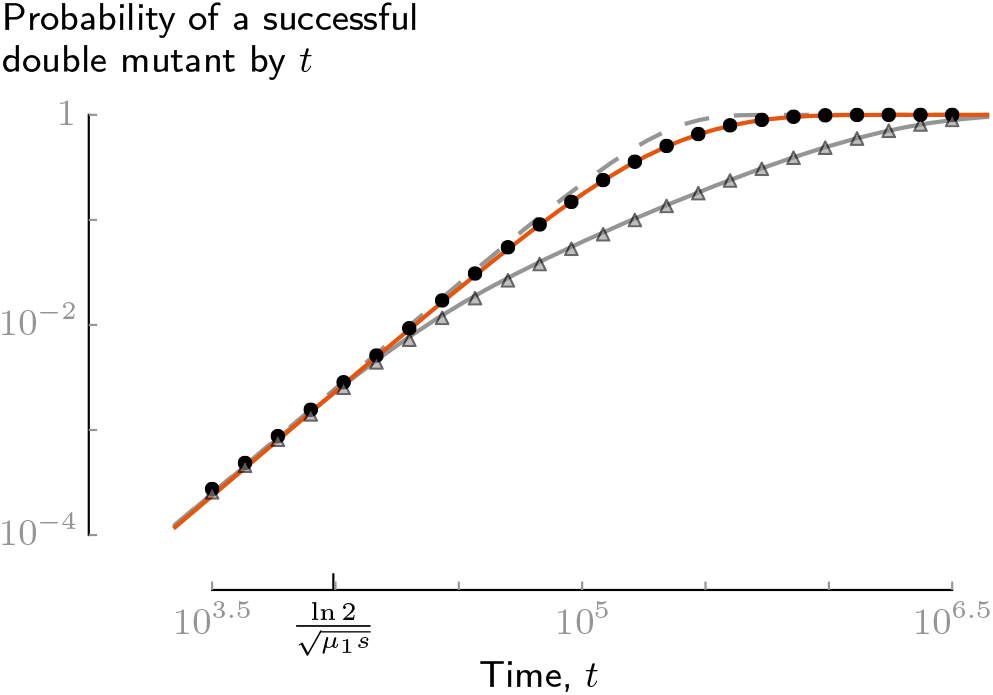
The probability 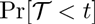 that a successful double mutant arises by time *t* is only increased by subdivision for times later than the average drift time in an unstructured population (here, approximately ln(2)/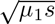. The grey line shows 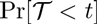 for an unstructured population, while grey triangles show estimates from simulations. The orange line shows 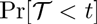 for a subdivided population of equal total size in which 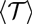 is approximately five times smaller, while black dots show estimates from simulations. The dashed grey line shows the prediction for 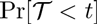 under neutral semi-deterministic dynamics. Here, single mutants are neutral for tunneling in the unstructured and subdivided populations. Parameters are *N*_T_ = 5×10^5^, *µ*_0_ = 10^−8^, *µ*_1_ = 10^−7^, *α* = 0.5, *δ* = 0, and *s* = 0.05; the subdivided population is split into *L* = 10^3^ demes of *N* = 500 individuals each, with migration rate *m* = 10^−5^.

For unstructured populations and subdivided populations with *δ* ≪ *α/N* that are in the NSD regime 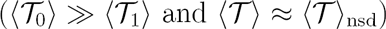, the probability of a successful double mutant by *t* is

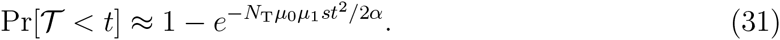

Equation (31) forms an upper bound on the probability of adaptation by time *t* regardless of the level of subdivision, which is approximately met for 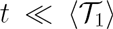. As the degree of subdivision is increased while *N*_T_ is held fixed, the average drift time increases and along with it the range of times for which 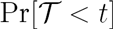 is given by (31), as seen in Figure 9.

### Subdivided populations with *δ* ≫ *α/N*

When single mutants are locally deleterious, we determine 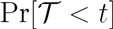 by separately considering the probabilities that a successful double mutant has been produced by a type-A lineage or by a type-B lineage before *t*. In general, due to their longer drift times, type-B lineages are less likely to contribute to 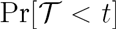 at very early times than at times 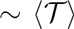. If 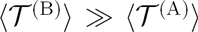, then type-A lineages form the dominate contribution to 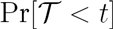 for all *t* and subdivision has no effect on the distribution. Therefore, we assume that *N*_T_*µ*_0_/*α* ≪ *ηδ*^2^/*µ*_1_*s* and *m* ≪ *α/N*, so that 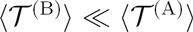 and adaptation typically occurs by type-B lineages. If 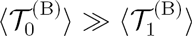, then 
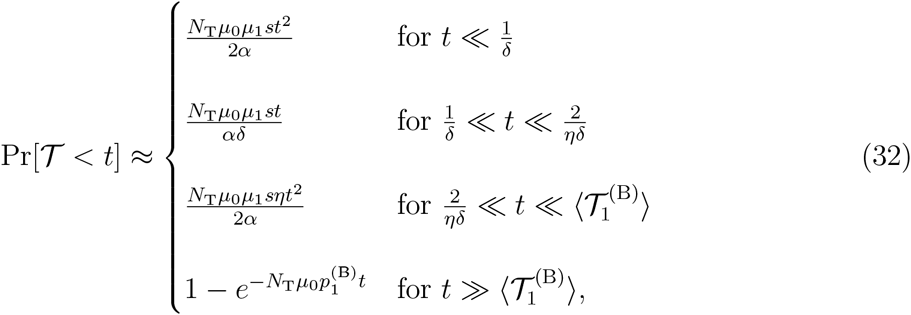
where 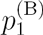 is given by (24) and 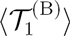 by (25). For *t* ≪ 2/*ηδ*, the probability of adaptation is determined solely from type-A lineages. The two regimes for *t* ≪ 2/*ηδ* correspond to whether *t* is shorter or longer than the average drift time ≈ 1/*δ* of a type-A lineage. For *t* ≪ 2/*ηδ*, the probability of adaptation is determined solely by type-B lineages. Again, the two regimes here correspond to *t* shorter or longer than 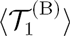. If instead 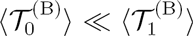, then 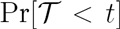 is given by (32) for *t* ≫ 2/*ηδ* and by 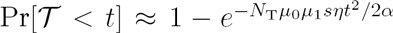 for t ≫ 2/*ηδ*. In both cases, type-B lineages only make the dominant contribution to 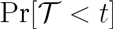 at times *t* ≫ 2/*ηδ*.

These results show that if *δ* ≫ *α/N*, the range of times over which subdivision has hardly any effect on 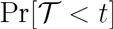 extends beyond the average drift time in an unstructured population, 1/*δ*, to the much longer time 2/*ηδ*. For 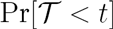 to be significantly increased by subdivision when *δ* ≫ *α/N*, it is necessary and sufficient both that 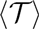 is significantly decreased and that *t* ≫ 2/*ηδ*.

## 7 Discussion

We have described the general ways in which population subdivision affects valley crossing in asexual species and given a complete analysis of valley-crossing dynamics for the island subdivision model. The competitive assortment caused by subdivision allows single-mutant lineages to have larger weights, with the result that single mutants have a greater probability *p*_1_ of being successful. Decreasing deme size or migration rate increases the level of assortment and thus *p*_1_, though significant increases in *p*_1_ occur only if demes and migration rates are small enough that successful single-mutant lineages fix in one or more demes before producing a successful double mutant. Subdivision also increases the drift and sweep times, which can greatly limit the extent to which subdivision will decrease the average time for the population to adapt or increase the probability that the population has adapted by a certain time.

In addition to the study of Komarova (2006), a number of additional studies on valley crossing in spatially structured asexual populations were published during the preparation of this manuscript. Komarova (2006) analyzed the cases of one neutral or deleterious intermediate in a population structured as a one-dimensional integer lattice, where competition occurs between individuals on neighboring sites on the lattice. Durrett and Moseley (2015) extended these results to include neutral intermediates on integer lattices of *d* ≥ 1 dimensions. Komarova *et al*. (2014) used simulations to consider one or more neutral, deleterious, or beneficial intermediates in a two-dimensional lattice, allowing for competition within a fixed radius. These lattice-population studies sought the waiting time for the first beneficial mutant to occur. Bitbol and Schwab (2014) considered the average time to fixation of the double mutant from a wild-type population in subdivision model very similar to ours, but only analyzed the limits of extremely frequent and extremely infrequent migration, while using simulations to consider variation in the migration rate. Spatial structure affects valley crossing by the same fundamental mechanism in all these studies (except in the second model of Komarova *et al*. (2014), for which spatial structure leads to lower total population sizes). Accordingly, all find that structure can accelerate valley crossing by increasing *p*_1_, although our results significantly extend these works; we compare our results with these works below. The seemingly contradictory finding of Takahasi (2007) that subdivision does not affect the crossing of a plateau in the absence of recombination is a consequence their unrealistic assumption that each mutation can occur only once in the population.

### Ability for subdivision to increase *p*_1_

In order for subdivision to significantly increase the probability that a single mutant is successful, we find the intuitive condition that successful single-mutant lineages must typically fix within one or more demes before producing a successful double mutant. This condition requires that both demes and migration rates are sufficiently small. Demes must contain fewer than *N*_×_ individuals to ensure that successful lineages do not simply tunnel within demes at low migration rates; this requirement was also found by Bitbol and Schwab (2014). In addition, we showed that for the island migration model, the migration rate *m* must be less than a threshold migration rate *ηα/N* for *p*_1_ to be significantly increased. If single mutants are locally neutral, the migration threshold simplifies to *α/N*, which corresponds to Wright’s classic result that large values of *F*_ST_ at neutral loci occur in the island model when *m* ≲ 1/*N*_e_ (Wright 1931). If single mutants are locally deleterious, the migration threshold is significantly less than *α/N*, reflecting the role of selection in limiting preventing locally deleterious lineages from obtaining large assortments by fixing within a deme (Whitlock 2002).

If *N* ≪ *N*_×_, then *p*_1_ increases with decreasing migration rates over a wide range. If double mutants are sufficiently beneficial (such that *s* ≫ *α/N*), then *p*_1_ increases until the maximal increase is approximately met when *m* ≪ *m*\*, corresponding to the isolated-demes regime in which single-mutant lineages that fix within a deme always produce a double-mutant deme. Bitbol and Schwab (2014) analyzed the isolated-demes regime and also estimated the migration boundary *m*\*. However, for *δ* ≪ *α/N*, the result in their Equation (10) (see also their Equation 17) overestimates *m*\* by a factor of ln(*L*) compared to the approximation *m*\* ≈ *Nµ*_1_*s/α* following from our (17). They estimate *m*\* by comparing the time for a single-mutant lineage that has fixed in a deme to produce a successful double mutant, versus the average time for the lineage to go extinct. However, the latter time is not relevant for single mutants that are neutral for tunneling, which is always the case for *m* near *m*\* when *δ* ≪ *α*/*N*. As a result, the approximation for *m*\* in Bitbol and Schwab (2014) overestimates the range of migration rates where *p*_1_ takes its limiting value as *m* → 0 when *δ* ≪ *α/N*.

### Consequences of the drift and sweep times

The effects that subdivision has on the drift and sweep times can greatly limit the ability for subdivision to accelerate valley crossing. Increases in the drift time caused by subdivision have two important consequences. First, subdivision can only decrease 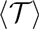 to a minimum of 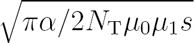, equal to the average waiting time for a successful double mutant under neutral semi-deterministic tunneling dynamics. If *N*_T_*µ*_0_/*α* ≫ max{1, *δ*^2^/*µ*_1_*s*}, then this minimum is met even in the absence of subdivision and subdivision has no effect on either the mean or the distribution of 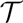. Komarova *et al*. (2014) observe a similar condition in their simulations for when structure does not affect 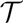 in two-dimensional lattice populations. Natural populations are frequently large enough that we might expect that *N*_T_*µ*_0_/*α* ≫ 1. This can often be expected for bacterial and viral populations, and can also occur in cancer-susceptible stem-cell populations (see discussion below). In these cases, significant decreases in 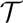 due to subdivision will only occur if single mutants are sufficiently deleterious.

The second important consequence of the drift time that subdivision can only increase 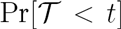 to a maximum of 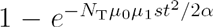. As a result, subdivision has no affect on 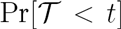 at times earlier than the average drift time in an unstructured population, even if it greatly reduces 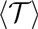.

These results for the minimum of 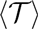 and maximum of 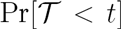 are both consequences of the fundamental trade-off between how spatial structure affects *p*_1_ and 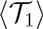. As a result, we expect them to apply in other types of spatial structure such as lattice population models. Other studies of tunneling have either only considered 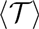 or assumed the exponential approximation approximation for 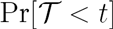 and so have failed to observe that the effects of subdivision on the average are in general not representative of the effects on the probability of adapting very quickly. This has critical implications for how the results of these studies should be interpreted when considering the effects of subdivision on 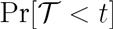 in applications to cancer development, which we discuss in further detail below.

Extreme subdivision also greatly increases the sweep time, which may more than offset any decreases in the time until the occurrence of a successful double mutant. Bitbol and Schwab (2014) also emphasized this point and found a similar expression for the average sweep time at low migration rates when *s* ≫ *α/N*.

### Other subdivision models

For many species, subdivision models where demes occupy geographic locations and individuals migrate preferentially to nearby demes are more realistic than the island model. Our results may also be used to predict valley-crossing dynamics in such cases, provided that demes have equal sizes and all individuals migrate at a fixed rate of *m* per generation. Our results for *p*_1_ and 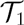 in the isolated-demes regime and for locally deleterious single mutants are approximately independent of the migration pattern between demes. Therefore, significant differences in our predictions are limited to regimes where *δ* ≪ *α/N* and *m* ≫ *m*\*, the regime in the island model described by the 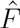 method. In general, single-mutant lineages that drift to ≫ *N* copies will experience larger assortments in models where migration is limited to nearby demes than in the island model. Thus, locally neutral single mutants will tend to have larger values of *p*_1_ outside of the isolated-demes regime, and may show significant increases in *p*_1_ at migration rates higher than *α/N*. Further study is required to determine how variation in deme size (e.g., if only a fraction of demes have fewer than *N*_×_ individuals) or variation in migration rates in different demes affect *p*_1_. We expect our results for the mean and distribution of 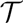 in terms of 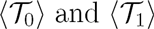 to continue to hold in these other subdivision models.

### Beneficial single mutants

When two mutations lead to a strongly beneficial complex adaptation, the first mutation may often have a smaller but still significant fitness advantage. We consider the effects of subdivision on the waiting time for such an adaptation in Supplementary Appendix H. Suppose that single mutants have a fitness 1 + *s*_1_ and double mutants have fitness 1 + *s* with 0 < *s*_1_ < *s*. If 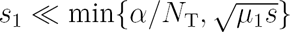, then single mutants are effectively neutral for the purposes of adaptation in both unstructured (Weissman *et al*. 2009) and subdivided populations, and subdivision can accelerate adaptation by increasing *p*_1_, On the other hand, if 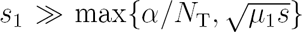, then in an unstructured population a successful single-mutant lineage will begin to sweep in the population before producing a successful double mutant. In such cases, subdivision can significantly increase the waiting time for a successful double mutant by slowing the sweep of the single mutant. Komarova *et al*. (2014) also observed this effect in simulations of two-dimensional lattice populations.

As with neutral and deleterious single mutants, extreme subdivision moves the mean and distribution of 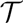 for beneficial single mutants towards the values predicted under neutral semi-deterministic tunneling. We consider the potential implications of this finding for cancer incidence rates below.

### Effects of subdivision in nature

To determine whether subdivision affects tunneling dynamics in nature, we must first ask whether natural populations are sufficiently subdivided that successful single-mutant lineages are likely to fix within a deme. Empirical measures of *F*_ST_ and related statistics using polymorphism data can provide some insight into the levels of competitive assortment in natural populations. Values of *F*_ST_ near 1 have been found in some species, such as the nematode *C. elegans* and in plant species with limited pollen dispersal (Charlesworth and Charlesworth 2010). Frost *et al*. (2001) found values of *F*_ST_ ranging between 0.08 and 0.59 for subpopulations of HIV in pulps within the spleens of infected patients. However, reported values of *F*_ST_ provide only limited insight into levels of competitive assortment, since they typically consider assortment over spatial scales much larger than the spatial scale at which individuals compete with one another. For example, studies commonly report estimates of *F*_ST_ between large geographic regions that most likely each consist of many smaller demes. Estimates of *F*_ST_ between regions likely underestimate competitive assortment, since rare mutant lineages are more likely to fix within a small deme than within a large geographic region.

Frequent local extinctions of demes followed by recolonization from a small number of founder individuals may be an important driver of large assortments in natural populations. For example, Frost *et al*. (2001) inferred that such dynamics drive the high values of *F*_ST_ in HIV populations within the spleen. Strong genetic hitchhiking also occurs in HIV (Zanini and Neher 2013; Zanini *et al*. 2015) and may act similarly by allowing a single-mutant lineage to quickly hitchhike to fixation on a linked beneficial mutation. Founder effects and hitchhiking may be particularly important in allowing locally deleterious lineages to reach fixation within a deme, but introduce additional complications and so further study of how these dynamics interact with subdivision to affect tunneling is needed.

Finally, we note that extreme subdivision occurs in two examples of tunneling in nature. The mitochondrial population of a mammalian species is divided into subpopulations corresponding to separate germlines, each with a small effective population size of *N*_e_ ≲ 200 (Howell *et al*. 1992; Jenuth *et al*. 1996) and no migration between them. These mitochondrial populations regularly cross valleys involving changes in the two complementary nucleotides that bind together to form the stems of tRNAs (Meer *et al*. 2010). In such cases, the first mutation alone is deleterious while the two mutations together are neutral. Meer *et al*. (2010) present genomic evidence that the deleterious single mutants remain at low frequency in the species’ population of mitochondria. Given the low germline effective population size and typical mitochondrial mutation rates, it is likely that the single-mutant lineage fixes within the germline before gaining the second mutation. Subdivision may increase the rate of tunneling by minimizing selection against single-mutant lineages within germlines. However, fixation within the germline is also thought to increase selection acting on mitochondrial mutations at the level of competition between hosts (Bergstrom and Pritchard 1998; Neiman and Taylor 2009), and so it is possible that subdivision may in fact reduces the rate of tunneling. Human tissues in which tunneling can give rise to cancer are also often extremely subdivided, and we now consider the implications of our results to this process.

### Applications to the development of cancer

Our results provide insight into how the structure of human tissue might influence the development of cancer, which we illustrate by considering the waiting time for a stem cell to gain a loss of function (LOF) mutation in both copies of a tumor suppressor gene (TSG). Details for our parameter estimates and calculations are given in Supplementary Appendix I. First hits to TSGs in healthy stem cells have been estimated to occur at rates of ~ 10^−6^ to ~ 10^−7^ per generation, while second hits are thought occur somewhat faster as loss-of-heterozygosity mutations may occur that replace the functional with the non-functional copy. We suppose that the second hit has a rate of *µ*_1_ = 10^−6^ per generation. Cellular reproduction within somatic tissues is similar to that in the Moran model, so that *α* ~ 1. If the first hit is neutral, then the average drift time if the population were unstructured is 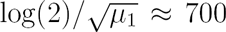 generations if double mutants always lead to cancer and longer if only a fraction do. The number of generations experienced by stem cells in cancer-susceptible tissues over an 80 year human lifespan ranges from a high of ~ 6000 in the colon to ≲ 80 for many tissues (Supplementary Appendix I). Thus, if single mutants are typically neutral then the average drift time without structure may be comparable to or longer than a typical human lifespan for many cancers, greatly limiting the ability for structure to increase the probability of acquiring two hits. Komarova (2006), Komarova *et al*. (2014), and Durrett and Moseley (2015) claim that spatial structure should increase the probability of cancer when the first hit is neutral, but ignore the drift time in their calculations and only account for the time until a successful single mutant appears.

In the colon, where a relatively large number of stem-cell generations occur, the susceptible stem-cell population is very large, at *N*_T_ ~ 8×10^7^ (Supplementary Appendix I). Colon cancer often begins by two hits to the APC gene, for which we estimate that *N*_T_*µ/α* ~ 50. Thus, if the first hit is neutral, we predict neutral semi-deterministic tunneling regardless of the level of subdivision. However, Vermeulen *et al*. (2013) recently showed that some single LOF mutants carry a large fitness advantage over wild-types within a crypt. This advantage is large enough that single mutants would likely sweep before producing a successful double mutant if the population were unstructured, but instead are confined to a single crypt due to the extreme subdivision present in the colon (Supplementary Appendix I). In reality, different LOF mutations are likely to have different fitness effects and cannot necessarily be lumped together into one mutation rate. However, if a large fraction of LOF mutations have a similar selective advantage, then subdivision in the colon likely greatly limits the appearance of stem cells with two LOF mutations.

### Conclusion

Population subdivision likely has important but complex effects on valley crossing in nature. Our results can be used to predict the effects of a broad range of different types of spatial structure on valley-crossing dynamics when recombination between mutations can be ignored. Our assumptions about population dynamics will frequently be invalid in nature due to processes we have ignored, such as extinction-recolonization dynamics, genetic hitchhiking, and selection at levels of population structure higher than demes. These require careful further consideration before we fully understand the effects of subdivision on tunneling in nature. However, we hope that our results may provide a foundation for studying these processes and for incorporating the effects of subdivision and spatial structure more generally in future theoretical and empirical investigations of valley crossing.

## Acknowledgements

I am grateful to I. Arbisser, B. J. Callahan, N. Creanza, M. D. Edge, M. W. Feldman, and D. B. Weissman for invaluable discussions and comments on the manuscript. I was supported during this research by the Stanford Department of Biology, the Stanford Genome Training Program (SGTP; NIH/NHGRI) and the Stanford Center for Computational, Evolutionary, and Human Genomics’ Fellows program. Computing resources were provided by NIH grant GM28016.

